# Spatio-temporal heterogeneity in hippocampal metabolism in control and epilepsy conditions

**DOI:** 10.1101/2020.06.17.155044

**Authors:** Giulio E. Brancati, Chahinaz Rawas, Antoine Ghestem, Christophe Bernard, Anton I. Ivanov

**Author notes:** ***Corresponding Author:*** Ivanov Anton, PhD, Institut de Neurosciences des Systèmes, Inserm U1106, Aix Marseille University, Faculté de Médecine de la Timone, 27 bd Jean Moulin, Marseille, 13005, France. Contributed equally to this work as first authors (Giulio E. Brancati, Chahinaz Rawas). Contributed equally to this work as senior authors (Christophe Bernard, Anton I. Ivanov).

## Abstract

The hippocampus’s dorsal and ventral parts are involved in different operative circuits, which functions vary in time during the night and day cycle. These functions are altered in epilepsy. Since energy production is tailored to function, we hypothesized that energy production would be space- and time-dependent in the hippocampus and that such organizing principle would be modified in epilepsy. Using metabolic imaging and metabolite sensing *ex vivo*, we show that the ventral hippocampus favors aerobic glycolysis over oxidative phosphorylation as compared to the dorsal part in the morning in control mice. In the afternoon, aerobic glycolysis is decreased, and oxidative phosphorylation increased. In the dorsal hippocampus, the metabolic activity varies less between these two times but is weaker than in the ventral. Thus, the energy metabolism is different along the dorsoventral axis and changes as a function of time in control mice. In an experimental model of epilepsy, we find a large alteration of such spatio-temporal organization. In addition to a general hypometabolic state, the dorsoventral difference disappears in the morning, when seizure probability is low. In the afternoon, when seizure probability is high, the aerobic glycolysis is enhanced in both parts, the increase being stronger in the ventral area. We suggest that energy metabolism is tailored to the functions performed by brain networks, which vary over time. In pathological conditions, the alterations of these general rules may contribute to network dysfunctions.

**SIGNIFICANCE STATEMENT:** The dorsal part of the hippocampus is involved in spatial learning and memory processes, while the ventral is implicated in motivational and emotional behavior. These functions change during the night and day cycle, and they are altered in epilepsy. Here we show that energy production (glycolysis versus oxidative phosphorylation) varies along the dorsoventral axis in a circadian manner *ex vivo* in control mice. These rules are altered in experimental epilepsy. Thus, energy production may be tailored to the function performed by hippocampal subdivisions and to the time of the day. Alterations in epilepsy may contribute to seizure generation and cognitive deficits.

## INTRODUCTION

Energy production in brain cells is assumed to be optimized to perform tasks or activities in a brain region-specific manner (1–3). Simultaneously, neuronal activity should be adapted to minimize the energy expenditure required for its fueling (4, 5). Several metabolic pathways are available for energy production, in particular glycolysis and oxidative phosphorylation (1). Whether specific metabolic pathways are preferably recruited in different brain areas in a task-dependent manner is not known. The hippocampus is an ideal region to test this hypothesis. The dorsal hippocampus (DH) is involved in learning and memory associated with navigation, exploration, and locomotion, whereas the ventral hippocampus (VH) is involved in motivational and emotional behavior (6–8). These functions are supported by very distinct anatomical (9, 10), morphological (11–13), molecular (14–19), and electrophysiological (12, 13, 20–23) properties of hippocampal cells. The hippocampus structure is also highly heterogeneous at the gene level, from its dorsal to its ventral tip (24, 25), which may serve as a substrate for different functional networks related to cognition and emotion to emerge (7, 26, 27). Given the hippocampus’s heterogeneity, from structure to function, along its dorsoventral axis, our first hypothesis is that energy production is different between the VH and the DH.

If energy production is tailored to a given structure-function relationship, we predicted that a change in energy production should accompany a change in the hippocampus’s functional state. Epilepsy is a particularly relevant situation to test this hypothesis. The different types of epilepsies are associated with numerous metabolic and bioenergetic alterations (28). Hypometabolism of epileptic regions is a common signature of mesial temporal lobe epilepsy (TLE) in humans and animal models, such as the pilocarpine model (29, 30). Importantly, in patients with TLE, only the temporal part (including the hippocampus) is epileptogenic. The temporal part corresponds to the ventral part in rodents, which has been identified as the epileptogenic region in the pilocarpine mode (31). Our second hypothesis is that any dorsoventral organization of energy metabolism found in control condition is altered in epilepsy.

Finally, hippocampal functions demonstrate circadian regulation (32), in particular place cell properties (33), long-term synaptic plasticity (34), as well as memory and learning processes (35, 36). Our third hypothesis is that energy production, specifically the respective contributions of oxidative phosphorylation and aerobic glycolysis (3), vary as a function of the time of the day in control and epilepsy.

To test these three hypotheses, we used an ex vivo approach to evoke an energy-demanding electrophysiological activity standardized for the three independent variables considered here: time, region (DH/VH), and perturbation (control/epilepsy). As a first step towards a better understanding of the time regulation of hippocampal metabolism, we considered two time points during the night/day cycle: Zeitgeber 3 (ZT3) and Zeitgeber 8 (ZT8), as they correspond to low and high seizure probability in the temporal lobe epilepsy model used. We found that the control DH and VH have distinct time-dependent metabolic signatures regarding glycolysis and oxidative phosphorylation. In experimental epilepsy, there is no more dissociation between DH and VH at ZT3, but the regional difference re-appears at ZT8.

## MATERIALS & METHODS

For a detailed description of methods, please refer to the Supplementary Information Methods section.

### Tissue slice preparation

Adult (2-4 months) FVB male mice were anesthetized with isoflurane before decapitation at ZT3 and ZT8 (i.e. 10:00 and 15:00, respectively). The brain was rapidly removed (< 30 s) and placed in ice-cold artificial cerebrospinal fluid (ACSF). One hemisphere, left or right in alternation, was used to prepare dorsal slices (DHS), whereas the other hemisphere was used to prepare ventral slices (VHS) as described in (11).

### Synaptic stimulation and field potential recordings

Slices were transferred to a dual perfusion recording chamber continuously superfused (6 ml/min) with ACSF contained 5mM of glucose, warmed to 30°C. Stimulation electrodes were placed in s*tratum radiatum* between the CA3 and CA2 areas. The extracellular local field potential (LFP) was recorded in *stratum pyramidale* of the CA1 area. Stimulation intensity was set to standardize responses (see SI Methods and Fig.S1).

### Oxygen, glucose, and lactate measurements

A Clark-style oxygen microelectrode (tip diameter 10 μm, polarization −0.8V; Unisense Ltd, Denmark) was used to measure slice tissue pO_2_. Tissue glucose and lactate concentrations were measured with enzymatic microelectrodes (sensing tip diameter 25 μm, length 0.5 mm, polarization 0.5V; Sarissa Biomedical, Coventry, UK). To avoid interaction between enzymatic glucose and lactate electrodes (see SI Methods), glucose and lactate measurements were done in separate sets of experiments.

### NAD(P)H and FAD^+^ fluorescence imaging

Slices were epi-illuminated with monochromatic light (pE-2 illuminator, CoolLed, UK), using 365 nm for NAD(P)H and 440nm for FAD^+^. The emitted light intensity changes were measured in *stratum radiatum* near the site of LFP recordings in the regions of interest ~1mm distant from the stimulation electrode tip using ImageJ software (NIH, USA). In the text, when referring to the optical signal, we use the term NAD(P)H (see explanation in SI Methods) while the abbreviation of reduced and oxidized forms of a coenzyme, NADH and NAD^+^, are employed when we discuss biochemical processes.

### Statistical analysis

To evaluate the difference between metabolic parameters values measured in dorsal and ventral slices we used in parallel two statistical approaches: the significance tests (parametric or non-parametric depending on the type of data distribution) and estimation statistic of bootstrap samples (37)(see SI Methods for more details). In the text, we reported the difference size as: unpaired or paired mean difference value [lower bound, upper bound of 95%CI]. In some cases, when the absolute values were of interest, we reported the corresponding parameter name and its mean value with 95% confidence interval obtained using the bootstrap procedure.

In all figures the p values are expressed as: *- p<0.05, **- p<0.01, ***- p<0.001.

## RESULTS

### The ventral hippocampus favors glycolysis over oxidative phosphorylation at ZT3

#### Metabolic imaging reveals dorso-ventral differences

Changes in NAD(P)H fluorescence can reflect the activity of both cytosolic and mitochondrial metabolic processes, which can be partly disentangled based on their time course (38), (39). Since the trains of stimulation produced a similar electrophysiological response in ventral and dorsal slices (dorsal: 3.8 ± 1.6 mV*s; ventral: 4.0 ± 1.6 mV*s, n=20; difference −0.1[−1.3, 1.1]mV*s, see also Fig.S2), we could test possible different metabolic responses between the DHS and VHS. Stimulations trains (10 s at 10 Hz) induced similar global dynamic changes in NAD(P)H fluorescence in both hippocampal regions (Fig.1Aa, left panel): at the onset of the stimulation (gray bar), the fluorescence intensity decreased rapidly (oxidative dip), before rising and overshooting until reaching a peak (reduction peak) after the end of the stimulation, preceding a slow return to baseline (dashed gray line). In our experimental conditions, the initial dip indicates that oxidative phosphorylation is dominating over cytosolic glycolysis, while during the rise to the peak, cytosolic glycolysis is dominating (39). The initial dip peaked within ~5-6 s after the train stimulation onset. Its mean amplitude was larger in VHS than DHS (0.7 [−1.1, −0.4]%, Fig.1Ab). Although the mean amplitude of the reduction peak was 0.9% larger in VHS than DHS, the 95% confidence interval included the 0 value, [−0.1, 2.1]%, preventing to reach a strong conclusion (Fig.1Ac, left panel). However, in response to a 30 s 10 Hz train, there was a significantly larger reduction peak in ventral slices (2.1 [1.2, 3.0]%, Fig.1Ac right panel).

**Figure 1.**
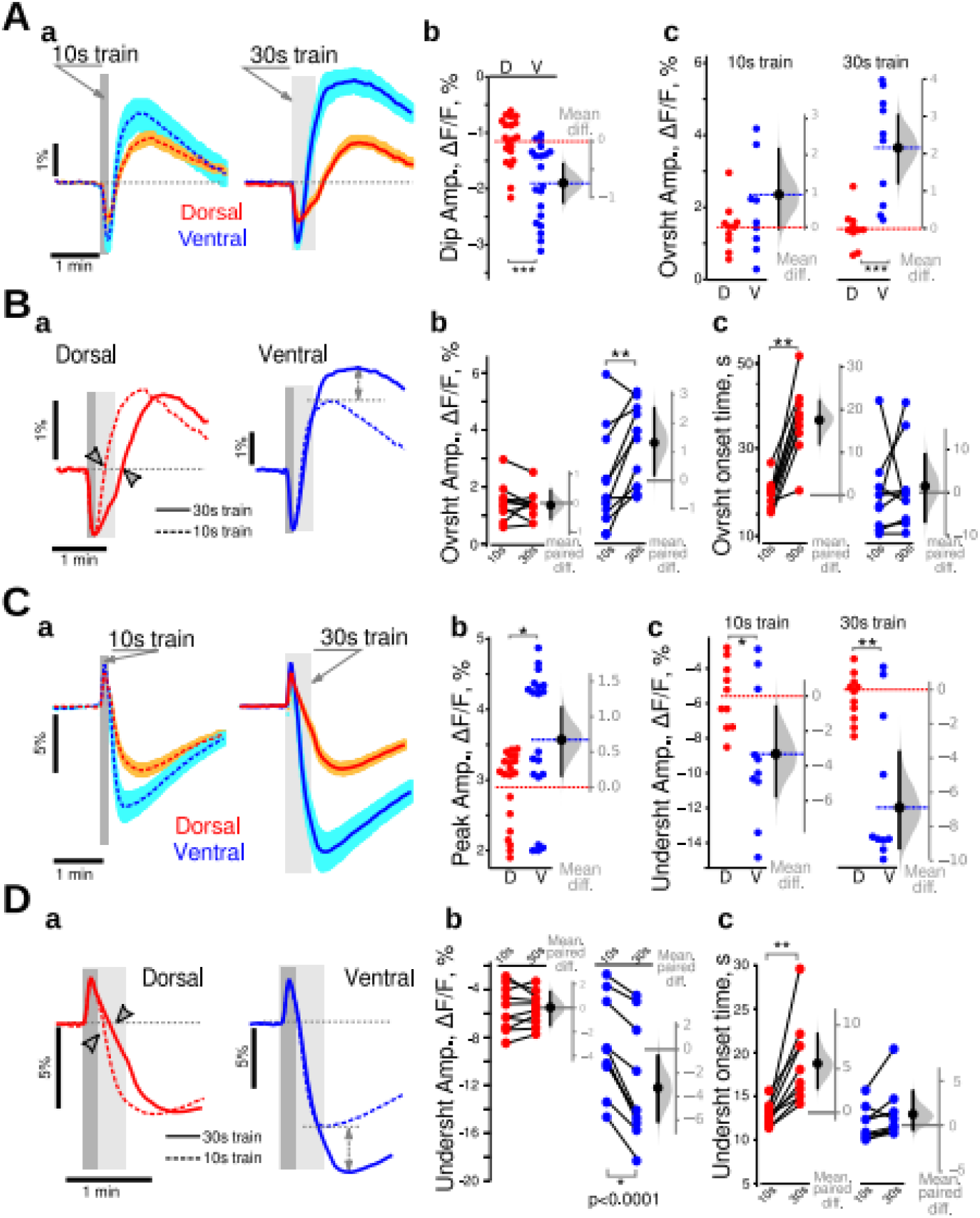
Dorso-ventral differences in energy metabolism revealed with imaging NAD(P)H and FAD^+^ in control animals at ZT3. **A.** Comparison of NAD(P)H fluorescence transients recorded in DHS (red) and VHS (blue). **a.** Averaged responses for 10 s (dark gray vertical bars) and 30 s (light gray vertical bars) trains. Both trains induced similar global dynamics, with an initial dip followed by an overshoot, and a return to baseline. Traces are averages from 10 independent experiments, with the light color shading indicating the SEM. (**b**) The amplitude of the initial dip was increased in the DHS (blue dots) compared to VHS (red dots). (**c**) Although not reaching statistical significance in the 10 s condition, 30 s long trains produced a larger overshoot in VHS than DHS. Metabolic activity is thus larger in ventral than in dorsal CA1. Experimental data are plotted on the left axis (**b, c**). The distribution (gray bell-shape) of the differences of mean values of bootstrap samples obtained by resampling of experimental data sets is plotted against the floating gray axis on the right, with the 0 point being aligned to the mean value measured in dorsal CA1 (red dashed line). The black circle with an error-bar shows the mean difference and its 95% confidence interval. Dashed red and blue lines correspond to the mean values of experimental data. **B.** Comparison of NAD(P)H transients evoked by 10 s (dashed line) and 30 s (continuous line) trains. **a.** In DHS (red lines), a longer stimulation delayed the overshoot onset (arrowheads) without affecting its amplitude. In VHS (blue lines), the overshoot onset times were identical, but the overshoot amplitude was larger with a 30 s train (dashed double-head arrows); **b,c.** Paired comparison of wave shape parameters of NAD(P)H transients evoked using 10 s and then 30 s long trains in dorsal (red dots) and ventral (blue dots) CA1. **C.** Comparison of FAD^+^ fluorescence transients recorded in DHS and VHS. **a.** synaptic stimulation with 10 s or 30 s trains produced similar global dynamics in FAD^+^ fluorescence, with a larger undershoot amplitude in VHS (blue lines) than DHS (red lines). Both initial peaks (**b**) and undershoot amplitudes (**c**) were larger in VHS. **D.** Comparison of NAD(P)H transients evoked by 10 s (dashed line) and 30 s (continuous line) trains. In dorsal CA1 (**a**, red lines), a longer stimulation delayed the undershoot onset (arrowheads on **a**, red dots on **c**) without affecting its amplitude (**b**, red dots). In VHS (**a**, blue lines) the undershoot onset times were identical in both cases (**c**, blue dots) but the amplitudes were larger for 30 s long trains (dashed double-head arrows on **a**, blue dots on **b**). On **Aa** and **Ca**, the averaged trace from 10 independent experiments is displayed—the light color shading indicating the SEM. Dark and light gray vertical bars show the times of 10 s and 30 s trains. Dark and light gray vertical bars indicate the time of 10 s and 30 s trains. Since the data set illustrated here passed Shapiro-Francia normality test, we used unpaired (**A, C**) and paired (**B, D**) t-test.

We then compared the differences between the two stimulation protocols (Fig.1B). Since the initial dip occurs within ~5-6 s, its amplitude was similar for both stimulations (Fig.1Ba). In the DHS, the 30 s train stimulation lengthened the oxidative dip resulting in a 18[12, 22]s delay in the onset of the overshoot (Fig.1Ba, red traces, gray arrowheads, and Fig.1Bc, red circles). However, the overshoot amplitudes remained unchanged(−0.1[−0.4, 0.2]%, Fig.1Bb: red circles). In VHS, the overshoot onset time was independent of the train duration (1.5[−6.7, 8.8]s, Fig.1Ba blue traces, Fig.1Bc blue circles), but the overshoot amplitude was significantly increased in the case of 30 s trains (1.3[0.1, 2.6]%, Fig.1Bb blue circles).

These results show that ventral CA1 is more active in terms of metabolism. When more energy is needed, VH enhances glycolysis (increased overshoot peak). In contrast, DH relies longer on oxidative phosphorylation (delayed overshoot).

Non-fluorescent FADH_2_ is highly concentrated in mitochondria. It is oxidized to fluorescent FAD^+^ in the respiratory chain and can be reduced again in the Krebs cycle to FADH_2_. FAD^+^ is also reduced at the glycerol-3-phosphate shuttle (G3PS) level as cytosolic NADH gets oxidized (see SI Discussion). At the onset of the stimulation, FAD^+^ fluorescence increased during 4 s (oxidative peak) then declined, reaching its minimum (undershoot) after the end of the train (Fig.1Ca). The FAD^+^ oxidative peak was larger in VHS than DHS (0.7[0.2, 1.1]%; Fig.1Cb). The undershoot amplitudes were also larger in ventral than dorsal slices for both 10 s(−3.3[−5.8, −0.6]%) and 30 s (−6.9[−9.3, −3.7]%) long stimulation trains (Fig.1Cc).

The paired comparison of FAD^+^ transients evoked by 10 s and 30 s long stimulation trains showed that the increased network activation did not affect the amplitude of the undershoot (0.1[−1.7,1.6]%) but delayed the onset time by 6[3, 10]s in DHS (Fig.1Da red traces, Db, Dc red circles). In contrast, in VHS, (Fig. 1Da blue traces, blue circles on Db and Dc), the undershoot amplitude was increased (−3.1 [−0.4, −6.1]%), while the undershoot onset time was not affected (1.2 [−0.5, 4]s). The FAD^+^ results thus support the NAD(P)H ones. Dorsal CA1 appears to rely more on oxidative phosphorylation than ventral CA1, which rapidly switches to glycolysis upon intense network activity. In ventral CA1, the large increase in cytosolic NAD(P)H will produce a large decrease in mitochondrial FAD^+^ via the G3PS, in addition to the decrease via the Krebs cycle, both processes regenerating FADH_2_.

In keeping with our hypothesis, the ventral and dorsal CA1 regions behave differently in terms of metabolism: ventral CA1 displays higher metabolic activity than dorsal CA1 and relies on different strategies to produce energy. To assess energy metabolism activity more directly, we measured the extracellular concentrations of lactate, glucose, and oxygen.

#### The ventral hippocampus is metabolically more active than the dorsal part in basal conditions at ZT3

We assessed energy consumption under basal conditions between VHS and DHS. In our experimental conditions, O_2_ and glucose concentrations are clamped by the ACSF, while lactate is not present in the ACSF. Figure 2A shows the depth profile of pO_2_ in slices. The depth with minimal pO_2_ value was denoted the ‘nadir.’ In the perfusion chamber, the ACSF pO_2_ was 520-650 Torr. The pO_2_ dropped to similar values at the slice’s surface (Fig.2A): 314±19 Torr in dorsal and to 274±18 Torr in the VHS (p=0.2, n=5). At the nadir (150-180 μm in both slice types), the pO_2_ was significantly lower in VHS than DHS (159±14 Torr vs 225±17 Torr, p=0.02, n=5, Mann-Whitney U test “M-W test”). Past the nadir, the pO_2_ began to increase as the slice’s bottom was also exposed to oxygenated ACSF in the dual perfusion chamber used here. Using two way ANOVA, we found that the type of slice accounts for 34% of the total variance (F_(1, 72)_=54.3, n=5, p<0.0001). When comparing pO_2_ values measured at a depth of 150-200 μm (Fig.2B, blue circles), we found that the concentration of oxygen was lower in VHS than DHS (−80[−140, −30]μM). There is thus a higher level of oxygen consumption in VHS in basal conditions.

**Figure 2.**
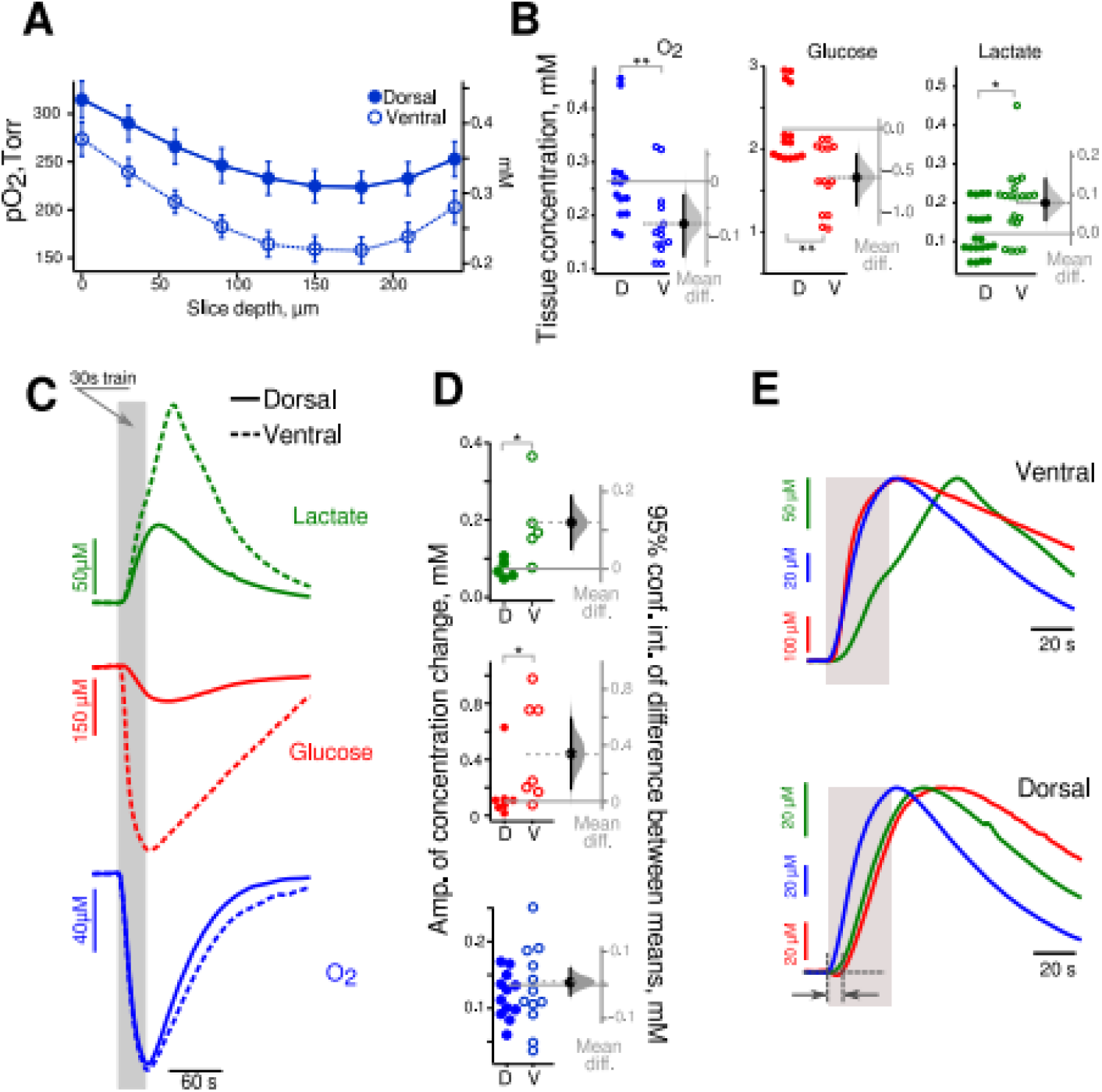
Increased aerobic glycolysis in the ventral hippocampus at ZT3. **A.** Mean extracellular oxygen content (pO_2_ on left axis, concentration on right axis) is lower in VHS (open circle) than in DHS (closed circles) slices at all depths suggesting higher oxygen consumption in basal conditions. **B.** Estimation plots of the mean difference of O_2_, glucose and lactate concentration at a 150-200 μm depth between VHS (open circle) and DHS (closed circles) slices. Metabolic activity is increased in ventral as compared to dorsal CA1 in resting (unstimulated) condition. **C.** averaged wave-shapes of the lactate (green), glucose (red), and oxygen (blue) responses to 30 s long trains (gray vertical bar) in dorsal (solid lines) and ventral (dashed lines) slices. **D.** Corresponding estimation of the mean differences in amplitude showing larger changes in lactate and glucose concentration in VHS than in DHS, whilst oxygen changes were similar. **E.** Averaged and normalized (to peak amplitude) time courses of glucose (red), oxygen (blue) consumption and lactate (green) release in VHS (upper panel) and DHS (middle panel). In VHS glucose and oxygen consumption started immediately at the onset of synaptic stimulation while in DHS glucose began to change 10 s later. Lactate changes had a similar time course in both types of slices.

Although glucose was clamped at 5 mM in the ACSF, its concentration was 2.3 mM in DHS and even smaller in VHS (−0.6[−0.9, −0.3]mM; Fig.2B, red circles). This finding is consistent with the difference in tissue pO_2_, because oxygen consumption is an indicator of the intensity of oxidative metabolism, which requires glucose. In parallel, we found more lactate (which can only come from cell metabolism in our conditions) in VHS than in DHS (0.08[0.04, 014]mM; Fig.2B, green circles), suggesting increased aerobic glycolysis. We conclude that there is a higher metabolic demand in ventral than in dorsal slices in basal conditions. We then assessed the same parameters when the network was stimulated.

#### Changes in extracellular glucose, lactate, oxygen in response to synaptic stimulation at ZT3

We used 30 s long stimulation trains as dorsoventral differences are better revealed in such conditions (Fig.1). Figure 2C shows that glucose (red) and oxygen (blue) levels decreased during the stimulation and started to recover after the train stopped. Lactate (green) changed in the opposite direction. Since lactate is not present in ACSF, its time course indicates production (via glycolysis), consumption (via oxidative metabolism), and washout by perfusion. Synaptic stimulation led to increased glucose consumption and higher lactate release in VHS than DHS (Fig.2C-D, 0.35[0.1, 0.6]mM and 0.12[0.07, 0.20]mM, respectively; M-W test p=0.03 for glucose and lactate, n=6). In contrast, oxygen consumption was identical (0.01[−0.02, 0.05]mM, n=12). These results suggest that aerobic glycolysis is increased in ventral compared to dorsal CA1, while oxidative metabolism is similar when the network is activated.

We then assessed the temporal profile of the responses. To facilitate the comparison, we normalized the traces to the peak amplitude and changed the polarity to make them positive-going (Fig.2E). Oxygen and glucose consumption had very similar time courses in VHS, while in DHS the glucose signal was delayed compared to oxygen (Fig.2E, bottom panel, dark grays arrows). In VHS, oxygen consumption started right after the onset of the synaptic stimulation with a very short time delay (mean delay 0.6[0.4, 0.9]s). Both signals reached their maximum simultaneously a few seconds after the end of the train (mean time to peak for oxygen: 4[2, 6]s; glucose: 6[2, 10]s). In DHS (Fig. 2E, bottom panel), oxygen consumption started immediately at the train onset (mean delay 0.4[0.1, 0.7]s) and peaked 3.2[2.1, 4.4]s after. Glucose time-course was slower: consumption started 6[3, 12]s after stimulation onset (M-W test: p=0.0003, n=7), and reached its maximum 23[12, 31]s later (M-W test: p=0.0003, n=7). These results suggest that oxidative metabolism (revealed by oxygen consumption) is recruited immediately in both parts of the hippocampus at the onset of the stimulation. Whereas oxygen and glucose consumption are temporally coupled in VHS, glucose consumption is delayed in DHS; suggesting the use of other energy sources during this delay. Alternative substrates include lactate, glycogen, amino acids, or intermediates of the citric acid cycle. Lactate is unlikely because we did not measure any decrease in lactate concentration at the stimulation onset (Fig.2E, green traces). On the contrary, in both VHS and DHS, lactate started to rise 3 s after stimulation onset (VHS vs DHS difference: 0.2[−1.5, 1.1]s). Alternate sources, in particular glycogen, will be addressed in the discussion.

We also compared the changes in metabolite concentration induced by 10 s and 30 s long trains (Table S1). We found that in both slice types, the longer stimulation increased the peak amplitude of oxygen identically. In contrast, the increase in glucose consumption was slightly larger in VHS than DHS (87 μM vs 56 μM, difference: 31[−21, 76]μM), while lactate release was more increased in VHS than DHS (91 μM vs 33 μM, difference: 49[10, 90]μM). This suggests enhanced aerobic glycolysis to fuel longer network activity ventral CA1.

Together, imaging and molecular sensing experiments reveal major differences between the hippocampus’s ventral and dorsal CA1 regions. The intensity of aerobic glycolysis is substantially higher in ventral CA1 in both resting and stimulated states, while dorsal CA1 relies more on oxidative phosphorylation. Thus, the coupling between energy metabolism and neuronal network activity changes along the dorsoventral axis (Fig.S3A).

### Metabolic alterations in experimental epilepsy

A stimulation, which intensity was similar to that used in control slices (260±30 μA, n=26), produced a 70-80% of maximal LFP response in pilocarpine injected (“pilo”) animals. The total integral values of the LFP responses to a 10Hz 30 s train stimulation were similar in dorsal and ventral slices (1.8±1.0 mV*s and 1.4±0.6 mV*s, respectively, difference 0.5[−0.6, 1.5]mV*s, n=13). These responses were 50% smaller than in control animals. Since we could normalize the electrophysiological responses in dorsal and ventral slices in pilo animals, we could directly compare metabolic responses. However, we could not compare directly metabolic responses between control and pilo mice because the electrophysiological responses could not be normalized between the two conditions.

#### Loss of regional metabolic NAD(P)H/FAD^+^ differences in epilepsy at ZT3

Optical NAD(P)H and FAD^+^ signals had similar general time courses in both VHS and DHS (Fig.3Aa, Ca). In contrast to control mice, the initial oxidative dip of NAD(P)H (Fig.3Ab) and oxidative peak of FAD^+^ (Fig.3Cb) were identical in VHS and DHS of pilo mice (Tab.S2). Similarly, the amplitudes of NAD(P)H overshoot and FAD^+^ undershoot were similar in VHS and DHS (Fig.3Ac, Cc, Tab.S2). Other major differences were apparent when comparing 10 s and 30 s long trains. Whereas the onsets of the NAD(P)H overshoot and the FAD^+^ undershoot were delayed in the DHS only in control animals, the delay was observed in both VHS and the DHS in pilo mice (Fig.3Bc, Dc, Tab.S2). Besides, the NAD(P)H overshoot amplitude was not increased anymore in VHS for 30 s long trains as compared to 10 s trains (Fig.3Bb, Tab.S2), while FAD^+^ undershoot amplitude was significantly increased in both types of slices (Fig.3Db, Tab.S2). These results show that, in epileptic conditions, the ventral and dorsal CA1 respond similarly at ZT3. Since responses in dorsal CA1 show a similar profile in control and epilepsy conditions (e.g. Fig.1Ba and Fig.1Da compared to Fig. 3Ba and Fig.3Da), we conclude that the VH adopts the metabolic signature of the DH. However, major differences appear as compared to the control condition. In particular, the increase in FAD^+^ undershoot while the NAD(P)H remains the same suggests an uncoupling between cytosolic and mitochondrial redox states in epilepsy.

**Figure 3.**
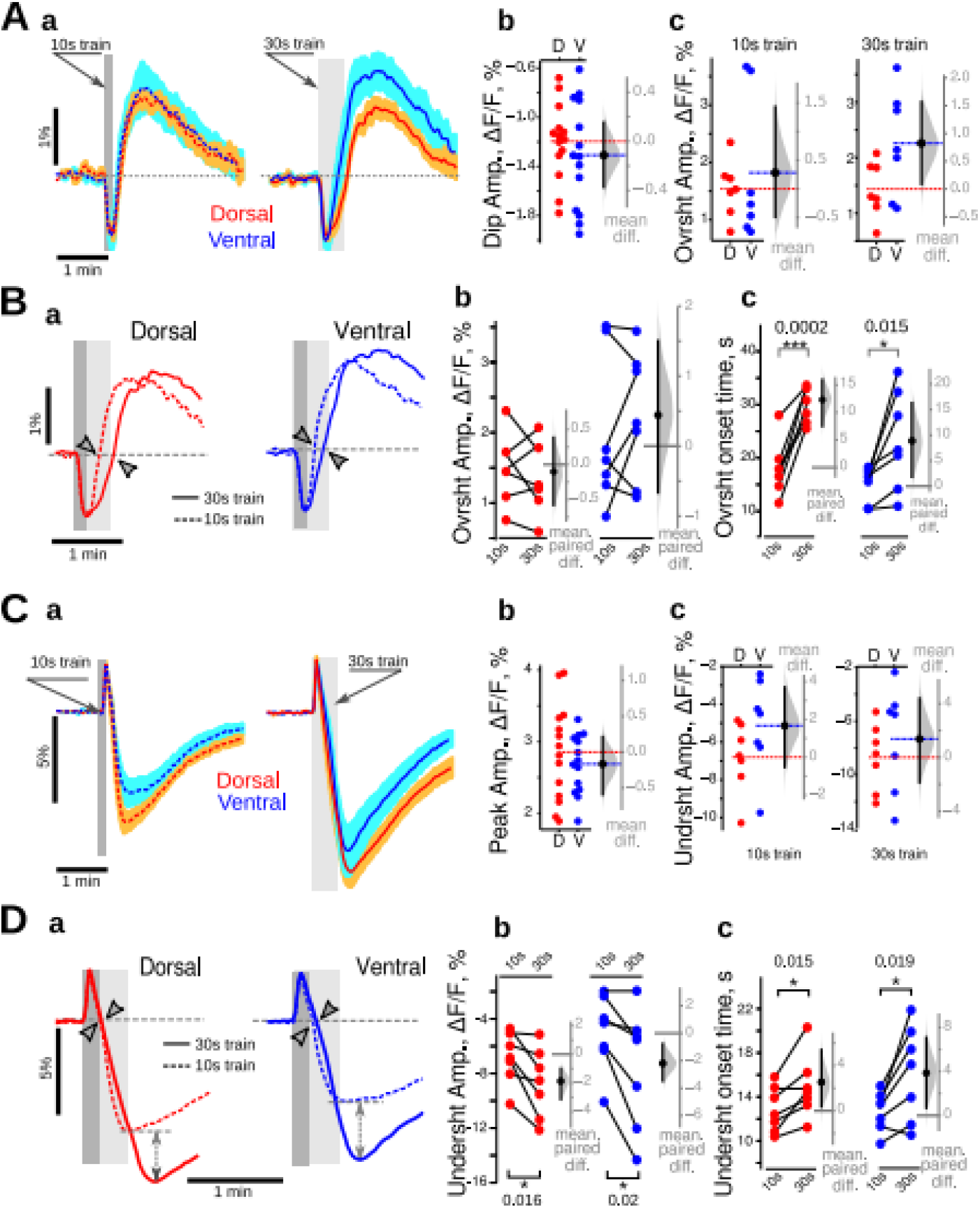
Loss of dorso-ventral differences in epilepsy at ZT3. **A. a.** The general time course of NAD(P)H was similar in DHS (red) and VHS (blue) in pilo mice for both 10 s (dashed lines) and 30 s (continuous lines) long trains. There was no difference in the amplitude of the oxidative dip (**b**) and overshoot (**c**) of NAD(P)H recorded in DHS (red circles) and VHS (blue circles). **B. a.** In both DHS (red) and VHS (blue), the 30 s long stimulation delayed the onset of the overshoot (**c**) without affecting its amplitude **(b**). **C. a.** The general time course of FAD^+^ was similar in DHS (red) and VHS (blue) in pilo mice for both 10 s (dashed lines) and 30 s (continuous lines) long trains. There was no difference in the amplitude of the oxidative peak (**b**) and undershoot (**c**) of FAD^+^ recorded in DHS (red circles) and VHS (blue circles). **D.** In both DHS (red) and VHS (blue), the 30 s long stimulation delayed the onset of the undershoot (**c**) and increased its amplitude (**b**). On **Aa** and **Ca**, the averaged trace from 7 independent experiments is displayed; for color code details, see Fig.1 legend. Since the data set illustrated here passed the Shapiro-Francia normality test, the unpaired (A, C) and paired (B, D) t-test were used.

#### Normalization of oxygen, glucose and lactate responses between ventral and dorsal CA1 in epilepsy at ZT3

We first estimated the basal levels of oxygen, glucose and lactate (Fig.4A). Although the lactate concentration was significantly lower in VHS than in DHS(−43[−60, −23]μM; M-W test: p= 0.001, n=11), the concentrations of glucose and oxygen were similar (glucose: −0.12[−0.56, 0.30]mM; oxygen: −0.04[−0.08, 0.01]mM). As for NAD(P)H and FAD^+^, the VHS seems to behave like the DHS in pilo mice in basal conditions. Although it was not possible to normalize the electrophysiological responses to stimulation trains to compare metabolic responses between control and epileptic animals, we could compare baseline concentrations (no stimulation condition). Since ventral temporal regions are part of the epileptogenic zone in the pilocarpine model, we hypothesized a large difference between control and epilepsy conditions in the VH. In DHS, the resting state concentrations of glucose and lactate in pilo mice were almost identical to those recorded in control animals (glucose: 0.3[−0.2; 0.6]mM; lactate: 14[−20; 40]μM), indicating no or very low effect of the epilepsy condition on the basal level glucose consumption and lactate release. In VHS, the difference between control and pilo condition was very large: glucose baseline concentration was 0.7 mM higher in pilo (95%CI [0.4; 1.1]mM; M-W test: p=0.0008), while lactate concentration was 110 μM lower (95%CI [−170; −8]μM, M-W test: p=0.002). This suggests a large decrease in aerobic glycolysis in the VH in epileptic animals in basal conditions at ZT3. Interestingly, oxidative metabolism was decreased similarly in DHS and VHS (pilo vs control difference in basal O_2_ concentration in dorsal: 100[40;140] μM; M-W test: p=0.0007; in ventral: 140[90; 190] μM, M-W test: p=0.0001). Thus, the altered metabolism in VH in epilepsy condition erases the regional difference between the DH and VH. This was further confirmed when analyzing the responses to stimulation trains (Fig.4B).

**Figure 4.**
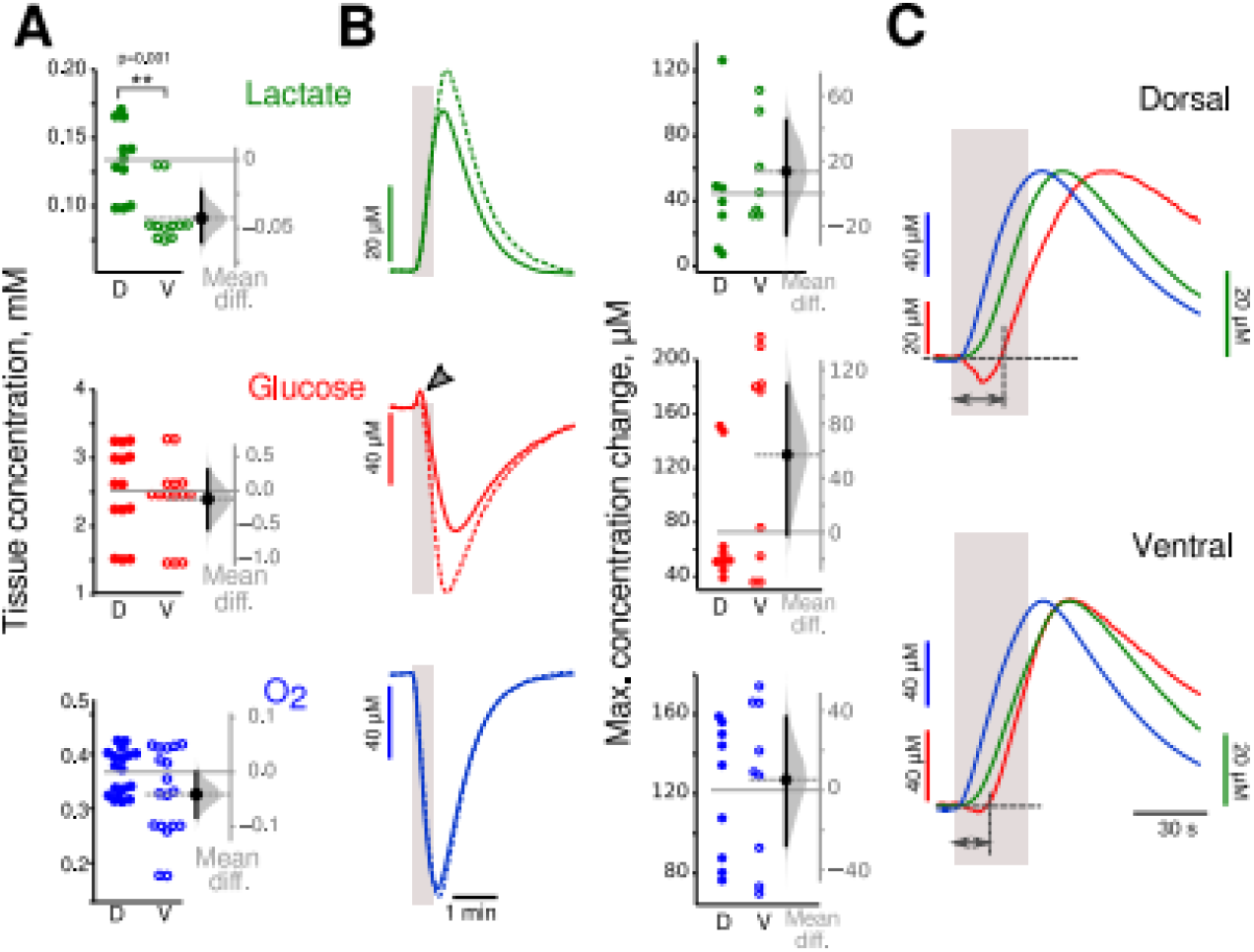
Changes in glucose dynamics in epilepsy at ZT3. **A.** Basal concentrations of glucose (red) and oxygen (blue) were similar in both slice types, while lactate (green) levels were significantly lower in VHS. **B.** The peaks of lactate, glucose, and O_2_ concentration in response to 30 s long trains were similar in DHS (solid line) and VHS (dashed line). Note that glucose concentration increased at the beginning of synaptic stimulation (gray arrowhead). **C.** Time course of averaged transient changes in lactate (green), glucose (red), and oxygen (blue). Traces are normalized to the peak, and polarity changed to make them positive going. In both DHS and VHS, glucose consumption started more than 10 s (double heads dark gray arrow) after the onset of synaptic stimulation (light gray bar). Since the data sets illustrated here did not pass the Shapiro-Francia normality test, we used the non-parametric M-W test.

The mean peak amplitudes of the three metabolites were similar in DHS and VHS (difference, lactate: 14[−25, 45]μM; glucose: 60[−2, 110]μM; oxygen 5[−28, 37]μM). Glucose consumption started more than 10 s after stimulation onset (mean delay 12[10, 14]s in dorsal, 10[6, 14]s in ventral, no difference between ventral and dorsal, see also Fig.4C, dark gray arrows). Interestingly, in 7 out of 9 DHS and 3 out of 9 VHS, we observed a positive deflection of glucose at the beginning of the synaptic stimulation (Fig.4B red traces, arrowhead), which interpretation will be addressed in the discussion. The delay in glucose consumption does not mean an increased latency of energy metabolism because the oxygen concentration (Fig.4C, blue trace) started to decrease immediately at the onset of electrical stimulation (mean delay 0.3[0.1, 0.8]s in dorsal and 1.0[0.7,1.5]s, no difference between dorsal and ventral) showing a good temporal coupling between electrical activity and oxygen consumption. Lactate production started to rise a few seconds (mean delay 6[5, 7]s in dorsal, 7[6, 8]s in ventral) after the stimulation onset (Fig.4C green trace) but before glucose consumption, suggesting that lactate production uses a substrate different from glucose (as it decreases later), such as glycogen (see Discussion section). Together these results confirm the disappearance of the regional difference between DH and VH in epileptic mice (Fig.S3C). Thus, at ZT3, there are important alterations of the metabolic rules in experimental epilepsy compared to control.

#### Differences between dorsal and ventral metabolism at ZT8 in experimental epilepsy

Since pilo mice have a higher probability to have a seizure at ZT8 than at ZT3 (see SI Methods), we assessed metabolic responses at ZT8 and found that dorsoventral differences appeared at ZT8. Neuronal network responses to the stimulation were similar in slices of both types (dorsal: 1.4 ± 0.9 mV*s, ventral 1.5 ± 0.8 mV*s, n=18, difference −0.1[−1.0, 0.8]mV*s) allowing direct comparison. The mean NAD(P)H overshoot amplitude was significantly larger (10 s train: 1.5[0.5, 2.4]%; 30 s train: 2.1[0.5, 3.2]%) while the oxidative dip was smaller(0.33[0.11, 0.51]%) in VHS than in DHS (Fig. 5A). FAD^+^ transients (Fig. 5C) were different only in their oxidative peak, which was smaller in VHS (mean difference −1.4[−1.9, −0.8]%). The mean undershoot amplitudes were similar for both 10 s (−0.7[−4.7, 2.9]%) and 30 s (−1.0[−5.1, 2.8]%) long trains. Both NAD(P)H and FAD^+^ signals recorded in DHS and VHS responded to the longer train with an increase of the overshoot/undershoot amplitude (Fig. 5B and D, double head dashed arrow). The onset time of FAD^+^ undershoot was delayed in DHS only by 2 s (Fig. 5, Dc), while the onset of NAD(P)H overshoot was not affected by the longer stimulation (Fig. 5, Bc). These results suggest that dorsal CA1 relies more on oxidative phosphorylation (larger amplitude of oxidative dip and peak of NAD(P)H and FAD^+^), while ventral CA1 preferentially uses aerobic glycolysis (larger NAD(P)H overshoot amplitude). However, to respond to energy demand induced by longer stimulation the slices of both types recruit aerobic glycolysis (increases in overshoot/ undershoot amplitudes). We conclude that dorso-ventral dissociation re-appears at ZT8 in pilo mice (Fig.S3D).

**Figure 5.**
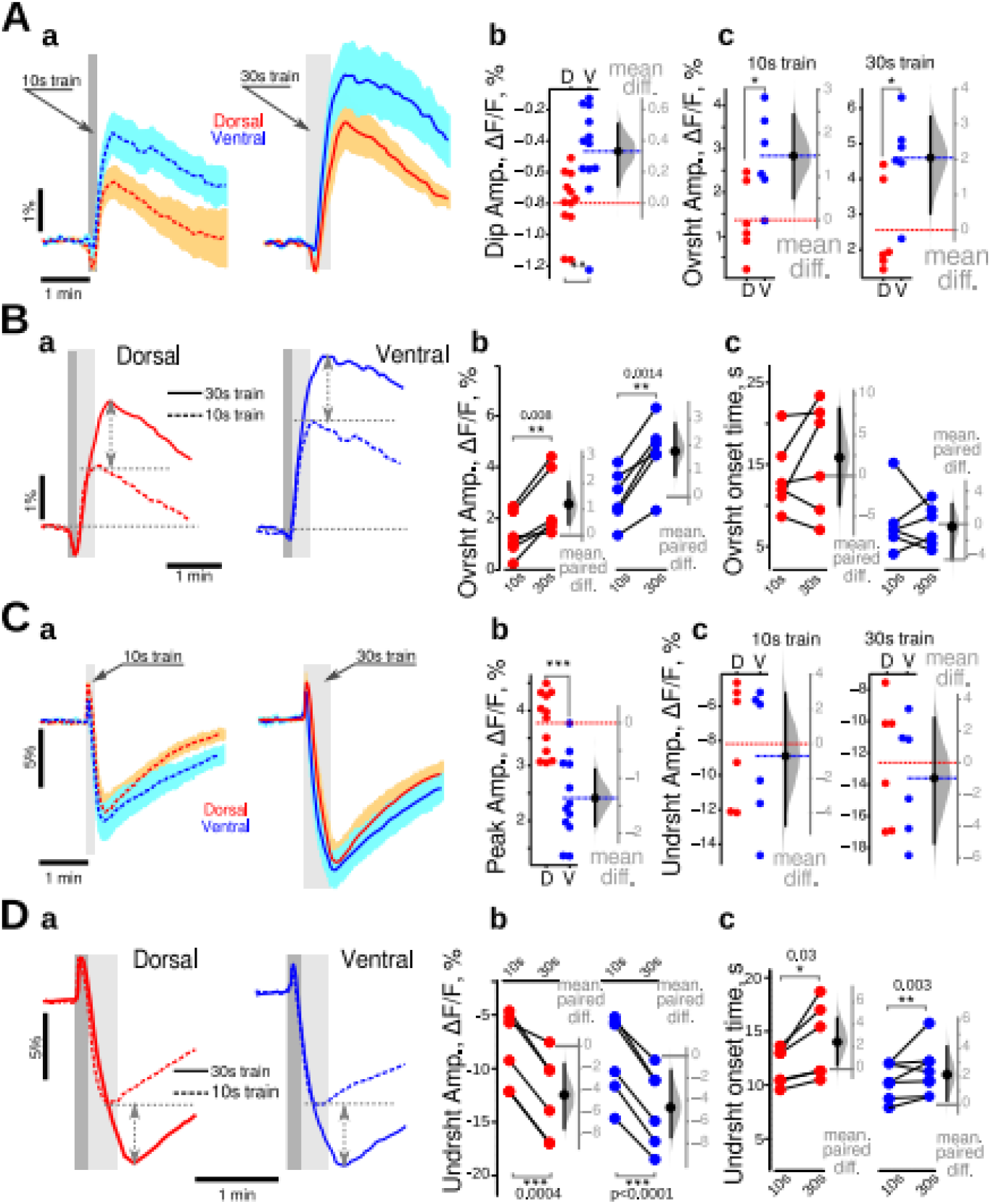
Dorsal-ventral differences in NAD(P)H and FAD^+^ responses in epileptic mice at ZT8. **A. a.** The general time course of NAD(P)H responses were similar in VHS and DHS. The oxidative peak of NAD(P)H fluorescence was larger in DHS than VHS (**b**), while the overshoot was larger in dorsal CA1 (**c**). **B.** Increasing the stimulation time increased the overshoot amplitude (**a**, double head arrows, **b**) without affecting its onset (**c**) in both regions. **C.** FAD^+^ signals were almost identical in DHS and VHS (**a**). The initial peak amplitude was increased in dorsal CA1 (**b**), while the amplitudes of the undershoot were comparable in both slice types independently from the train duration (**c**). **D.** Increasing the stimulation time increased the amplitude of the FAD^+^ undershoot (**a**, double head arrows, **b**) with only a minor effect on its onset time (**c**). **Aa** and **Ca** are averaged from 6 independent experiments; for color code details see Fig.1 legend. Since the data passed Shapiro-Francia normality test, we used the unpaired (A, C) and paired (B, D) t-test.

#### Increased aerobic glycolysis in ventral CA1 in epileptic mice at ZT8

In non-stimulated conditions, the extracellular concentration of lactate was similar between DHS and VHS (Fig.6A, difference 10[−20, 30]μM), in contrast to ZT3 (Fig.4A). However, the concentration of oxygen and glucose (Fig.6A) were significantly lower in VHS (glucose: −0.4[−0.70, −0.05]mM, oxygen: −0.15[−0.20, −0.09]mM). Thus, in basal conditions, VHS is more active with enhanced oxidative metabolism at ZT8, in contrast to the lack of differences at ZT3 (Fig.4). In response to electrical stimulation, we found an increased production of lactate (Fig.6B, 50[10, 80]μM) and increased consumption of glucose(0.23[0.05, 0.40]mM) in VHS, while oxygen consumption was similar(−10[−30, 15]μM). These results are consistent with the NAD(P)H ones, revealing enhanced aerobic glycolysis in VHS (larger overshoot and larger changes in lactate and glucose).

**Figure 6.**
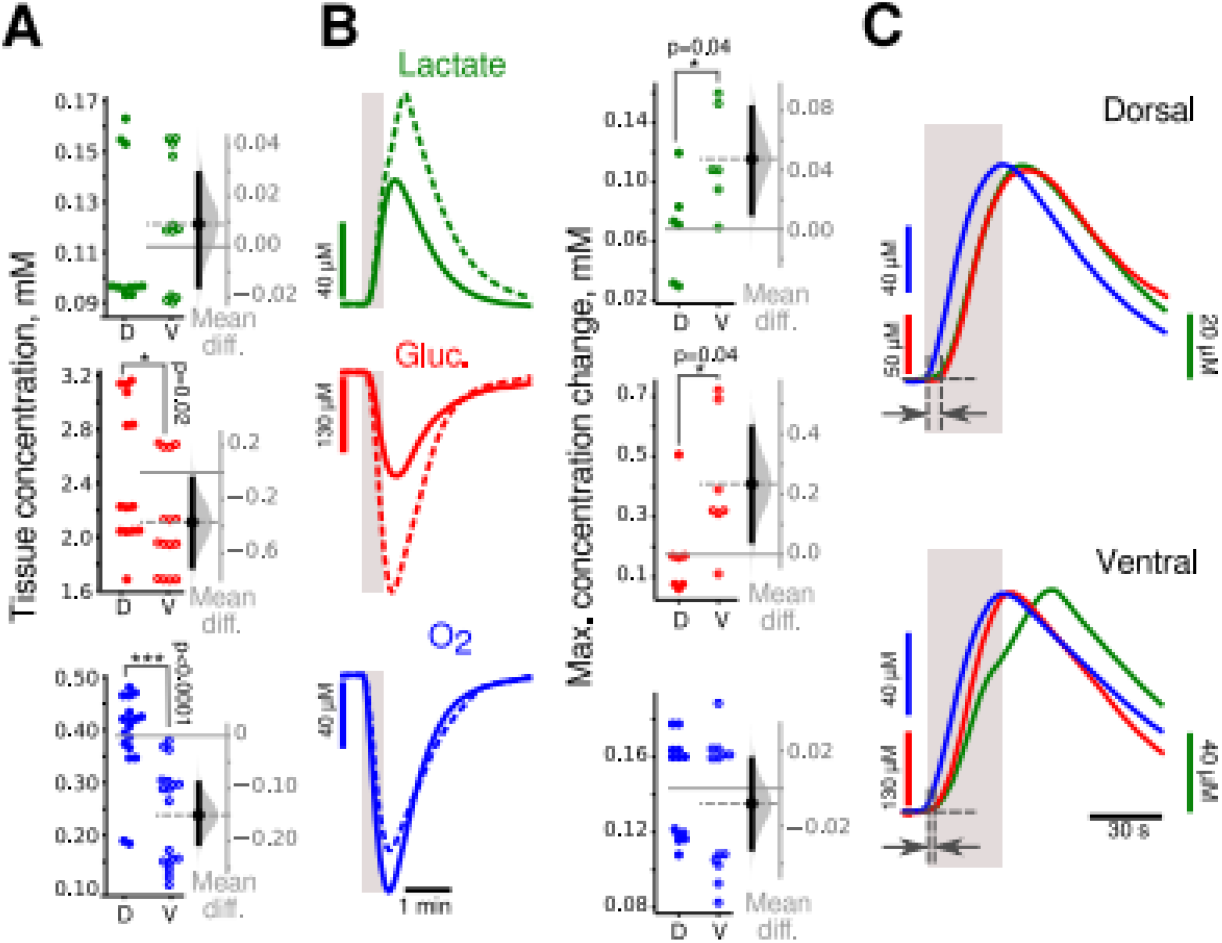
Increased aerobic glycolysis in ventral CA1 in epileptic mice at ZT8. **A.** In basal conditions, tissue concentrations of glucose (red circles) and oxygen (blue circles) were lower in VHS suggesting that they consumed more glucose and oxygen that DHS. There was no detectable regional difference in lactate content (green circles). **B.** In VHS, synaptic stimulation (gray vertical bars) was associated with a larger glucose consumption (red dashed lines and open circles) and lactate release (green dashed lines and open circles) than in DHS (solid lines and closed circles), while oxygen changes were similar(blue lines and circles). **C.** Time course of averaged transient changes in lactate (green), glucose (red), and oxygen (blue). Traces are normalized to the peak, and polarity changed to make them positive going. In both slice types, glucose consumption started later (dark gray arrows) than oxygen, which onset was synchronous to the beginning of the electrical stimulation (vertical gray bar). Since the data did not pass the Shapiro-Francia normality test, we used the non-parametric M-W test.

As observed at ZT3, glucose consumption started with a delay (Fig.6C dark gray arrows). This delay was shorter in VHS than DHS (−3.0[−5.0, 0.5] s; M-W test: p=0.04, n=7), suggesting a preferential utilization of extracellular glucose rather than alternative energy substrate (see Discussion). There was no initial rise in glucose concentration in slices of both types as observed at ZT3 (Fig.4B red trace, arrowhead), indicating an improved energetic state (see Discussion).

Together, these results suggest a strong circadian alteration in the rules governing energy metabolism along the dorsoventral axis in epileptic mice. We thus directly compared the results obtained at ZT3 and ZT8 (Fig.7).

### Circadian alterations in energy metabolism in epilepsy condition

The NAD(P)H responses were substantially different between ZT3 and ZT8 in ventral and dorsal CA1 in pilo mice (Fig.7Aa dorsal-warm colors, ventral-cold colors): the initial dip was smaller, the overshoot larger, and started earlier at ZT8. Since the data followed a normal distribution (Shapiro-Francia normality test), we performed a variability analysis using a two-way ANOVA (Fig.7 Ab, Ac, Ad). The analysis confirmed that time of the day (ZT3 vs. ZT8) was extremely significant for all parameters describing NAD(P)H transients (p<0.001 for each parameter), and that there was no interaction between the time of the day and the type of slice (dorsal/ventral). The Bonferroni corrected post-hoc test did not confirm the time dependence of overshoot amplitude in dorsal slice (Fig.7Ac). The time-dependence of FAD^+^ signals was less apparent (Fig.7Ba) but still significant. Variability analysis showed that both undershoot parameters, amplitude and onset time, were influenced by the time of the day (two-way ANOVA, p=0.0014 for the amplitude, p=0.0002 for the onset time), while the oxidative peak was not influenced (p=0.06). The Bonferroni corrected post-hoc tests revealed that only the undershoot was larger (p<0.01) and started earlier (p<0.001) at ZT8 in ventral slices (Fig.7 Bc, Bd). These results suggest that aerobic glycolysis is enhanced (larger overshoot/undershoot amplitudes) and starts earlier at ZT8 than at ZT3, especially in VHS. However, NAD(P)H and FAD^+^ results do not strongly support a possible circadian regulation of oxidative metabolism since NAD(P)H oxidative dips were smaller at ZT8 (Fig. 7Ab) while FAD^+^ peaks were either larger in DHS or unchanged in VHS (Fig.7Bb). Comparing the oxygen, glucose, and lactate levels provided the necessary information to interpret the results. Oxygen consumption in stimulation conditions was similar at ZT3 and ZT8 (Tab.S3 and Fig.7Ca), while the decrease in glucose concentration was substantially larger at ZT8 than at ZT3 (Tab.S3 and Fig.7Cb top). VHS also released more lactate at ZT8, whereas in DHS this parameter was not affected (Tab.S4 and Fig.7Cc). Importantly, glucose consumption started early at ZT8 (Tab.S4 and Fig.7Cb bottom) in keeping with the facts that the onset times of NAD(P)H overshoots and FAD^+^ undershoots occurred earlier. An early glycolysis onset would advance NAD^+^ reduction, thus masking the oxidative dip, making its amplitude smaller.

**Figure 7.**
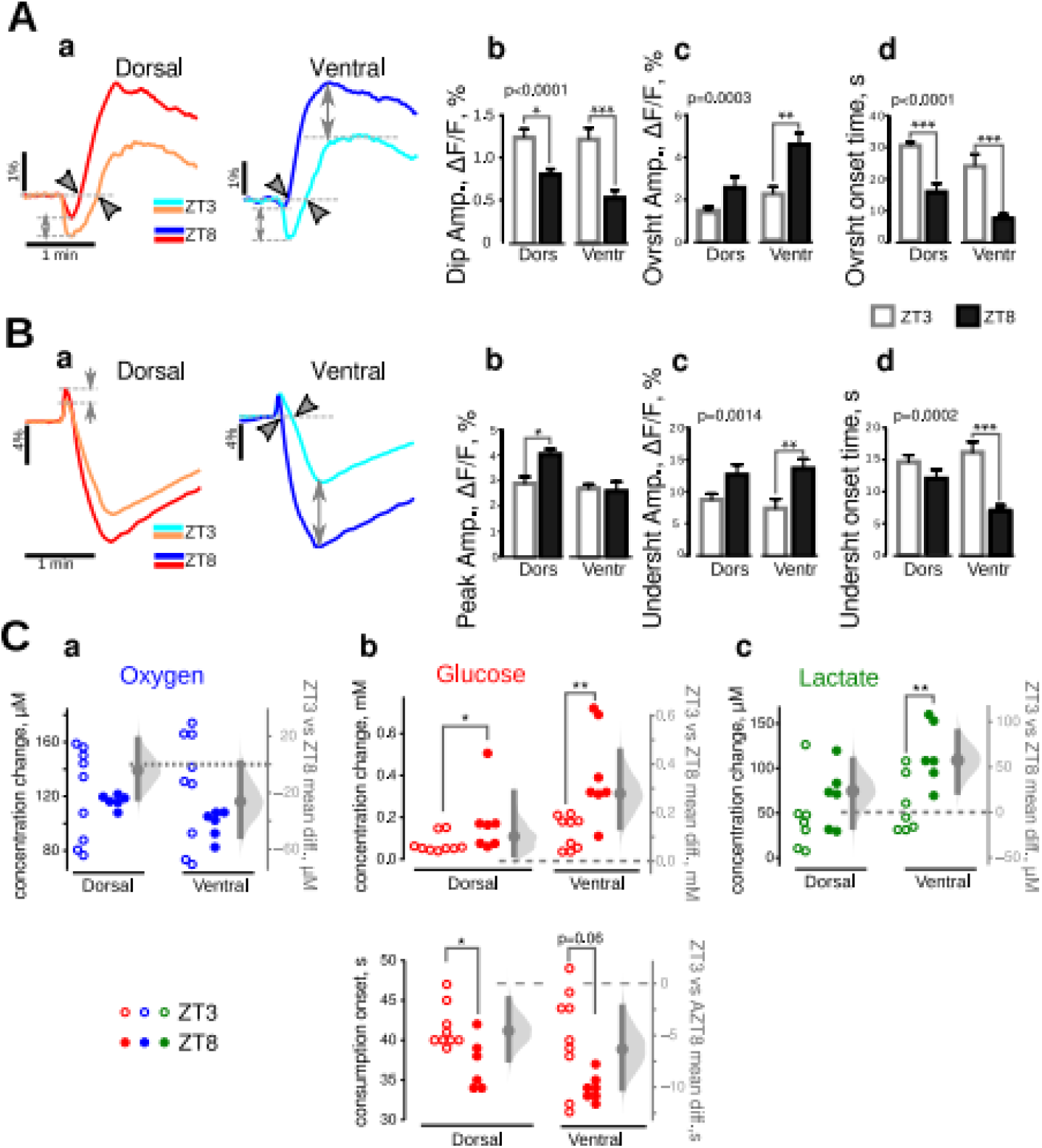
Accelerated and increased aerobic glycolysis at ZT8 in epileptic mice. **A. a.** Comparison of averaged NAD(P)H responses at ZT3 (light color traces) and ZT8 (dark color traces). Two way ANOVA revealed that all parameters describing the wave-shape of NAD(P)H transients (**a**-initial dip amplitude, **c**-overshoot amplitude, **d**-overshoot onset time) depend upon the time of the day (p values indicated in the top left corners of **b**,**c**,**d** panels), for both VHS and DHS (asterisks show the results of Bonferroni corrected post-hoc t-tests). **B.** FAD^+^ signals were also time of the day-dependent, but the effect was different between DHS (warm color traces) and VHS (cold color traces). In particular, the oxidative peak was different in VHS only (**a**-gray arrows). Two way ANOVA confirmed that the amplitude of undershoot (**c**) as well as its onset time (**d**) changed as a function of the time of the day (p values in the top left corners of **c**,**d** panels). Post-hoc tests indicated that this variability was significant mainly in VHS (asterisks). **C.** Oxygen (**a**), glucose (**b**), and lactate (**c**) changes induced by synaptic stimulation in DHS and VHS prepared at ZT3 (open circles) and ZT8 (closed circles). Although oxygen consumption did not change at ZT3 and ZT8, glucose concentration was larger (**b**, top panel) and started earlier (**b**, bottom panel) in slices made at ZT8. (**c**) The release of lactate in VHS was increased at ZT8 as compared to ZT3. Since the data in **A** and **B** were normally distributed (Shapiro-Francia normality test), we used the unpaired Bonferroni corrected post-hoc ttest. Since experimental data on **C** were not normally distributed, we used the M-W test.

We conclude that aerobic glycolysis is accelerated and enhanced at ZT8 compared to ZT3, in both ventral and dorsal CA1, while the intensity of oxidative metabolism is similar at both time points (Fig.S3C, D). Since metabolic properties are different between ZT8 and ZT3 in epilepsy conditions, we assessed whether metabolic properties are also time-dependent in control animals.

### Circadian regulation of metabolism in CA1 in control animals

The comparison of dorsal and ventral slices obtained at ZT8 led to the same conclusion as those prepared at ZT3 in control animals. The intensity of aerobic glycolysis was substantially higher in ventral CA1 in both resting and stimulated states (stimulation intensity 210-280 μV, LFP integrals 4.3 ± 1.8 mV*s in dorsal, 4.0 ± 1.5 in ventral, difference −0.4[−0.8, 1.1]mV*s n=12), while dorsal CA1 relies more on oxidative phosphorylation (Fig.8 and Tab.S4 for statistical analysis). The dorsoventral dissociation is thus maintained at ZT8 in controls.

**Figure 8.**
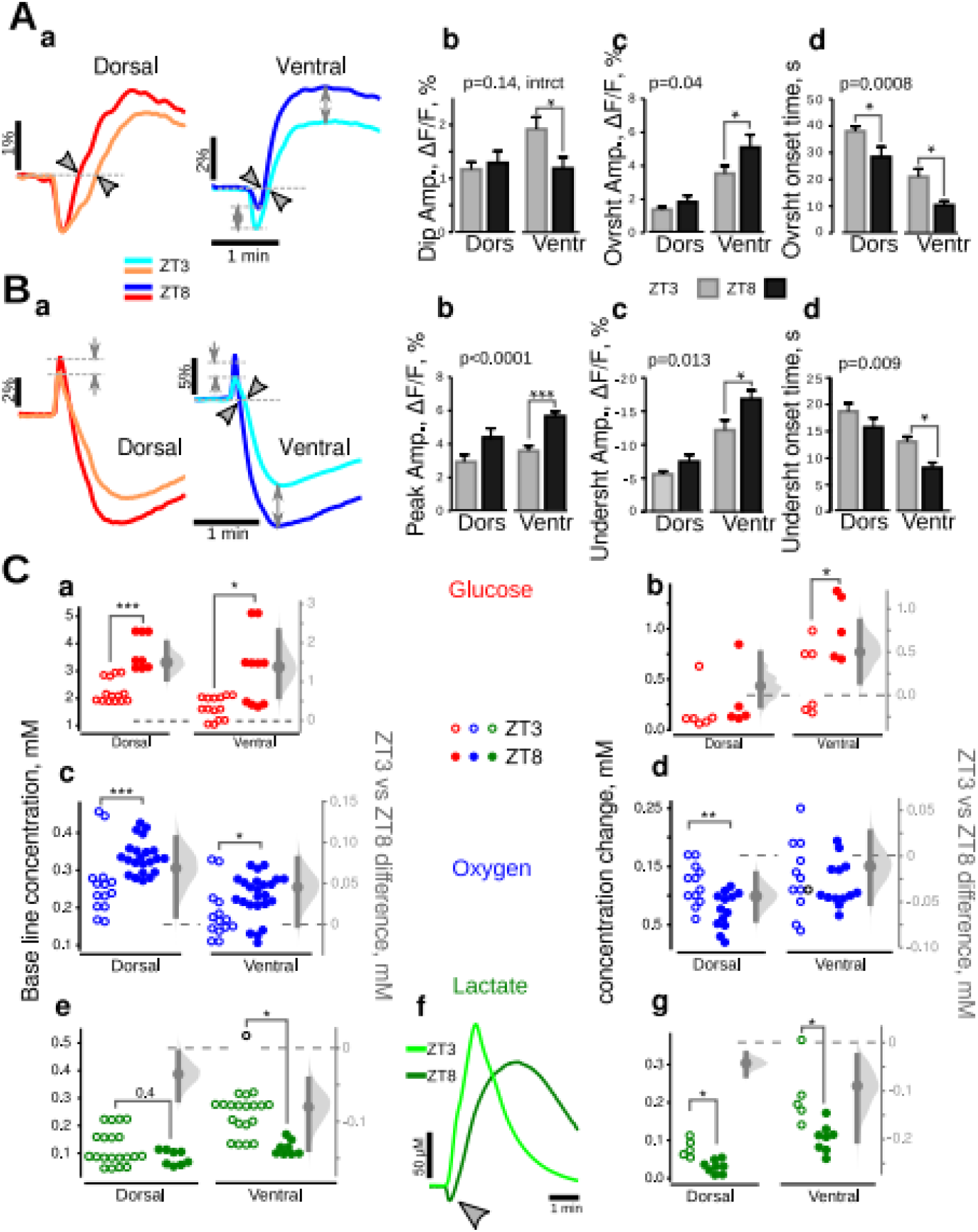
Dorso-ventral and temporal dissociations in metabolism in control mice. **A. a.** Averaged traces of NAD(P)H transients in DHS and VHS at ZT3 (light color traces) and ZT8 (dark color traces). (**b**) A two-way ANOVA showed an interaction between two factors, slice type and experimenting time, for dip amplitude, i.e. a different time regulation for DHS and VHS. The amplitude of overshoot (**c**) as well as its onset time (**d**) varied significantly with time in both slice types (p values in the top left corners of **c**,**d** panels). The Bonferroni corrected post-hoc t-test revealed that all three parameters describing NAD(P)H transients recorded in ventral slices varied with time while only the overshoot onset was modified in dorsal (**b**,**c**,**d** asterisks). **B. a.** Averaged traces of FAD^+^ transients show a higher oxidative peak, as well as a larger and earlier undershoot at ZT8 (dark color traces) than at ZT3 (light color traces). Two-way ANOVA confirmed that all these three parameters varied with ZT (p values in the top left corners of **b**, **c**, **d** panels) but, this variability was significant in ventral slices only (asterisks). **C.** Comparison of glucose, oxygen, and lactate concentration in quiescent and stimulated slices made at ZT8 (closed circles) and ZT3 (open circles). In both types of slices, the resting state glucose (**a**) and oxygen (**c**) concentrations were larger at ZT8 suggesting a lower level of metabolic activity that was consistent with the lower lactate concentration (**e**). Under stimulation conditions, glucose consumption did not vary (**b**) in dorsal slices. However, oxygen consumption (**d**) and lactate release (**g**) were lower at ZT8. In VHS, glucose consumption (**b**) was increased while lactate release (**g**) was decreased, probably due to lactate consumption as revealed with the concentration dip observed at the beginning of synaptic stimulation (**f**, gray arrowhead). Since experimental data on **C** were not normally distributed, we used the M-W test.

Clear differences appeared between DHS and VHS, when we compared the metabolic responses obtained at ZT3 and ZT8. In DHS, the parameters characterizing NAD(P)H and FAD^+^ responses (Fig.8 A and B) were similar at ZT3 and ZT8, except for a shorter overshoot onset time (Fig.8Ac).

The steady-state concentration of metabolites showed a decreased metabolism at ZT8: the levels of glucose (Fig.8Ca) and oxygen (Fig.8Cc) were higher (lower consumption), while lactate (Fig.8Cf) was lower. In stimulation condition, although glucose utilization was independent of time (Fig.8Cb), oxygen consumption (Fig.8Cd) and lactate release (Fig.8Cf, Cg) were smaller at ZT8. There is thus a weaker metabolic activity in DHS at ZT8 than at ZT3.

The situation was very different in VHS, in which both oxidative phosphorylation (FAD^+^ peak) and aerobic glycolysis (NAD(P)H overshoot amplitude and onset, FAD^+^ undershoot amplitude and onset) were enhanced at ZT8 as compared to ZT3 (Fig.8 A and B dark color traces vs. light colors).

Interestingly, the steady-state concentration of metabolites showed a decreased metabolism at ZT8 as for DHS: the levels of glucose (Fig.8Ca) and oxygen (Fig.8Cc) were higher (lower consumption) while lactate (Fig.8Ce) was lower.

In response to synaptic stimulation, VHS consumed oxygen similarly at both time points (Fig.8Cd) but absorbed more glucose (Fig.8Cb) and released substantially less lactate (Fig.8Cg) at ZT8 than at ZT3. Moreover, in 4 of 5 experiments, lactate concentration decreased at the onset of synaptic stimulation (Fig.8Cf, gray arrowhead). Since glucose consumption onset was not delayed compared to ZT3 (Tab.S4), the observed lactate dip may indicate that lactate consumption dominated over its production. We can rule out a downregulation of lactate production because it would decrease the amplitude of NAD(P)H overshoot; while we observed the reverse (Fig.8Aa, c). When consumed, lactate is transformed to pyruvate, which requires NAD^+^ reduction. The resulting NADH production, occurring at the very start of the stimulation, could mask the increase of the initial NAD(P)H dip (Fig.8 Aa, Ab). Therefore, since the FAD^+^ response clearly showed increased oxidative phosphorylation at the beginning of the response (Fig.8Ba, Bb), we conclude that this metabolic pathway was increased while aerobic glycolysis is downregulated at ZT8 as compared to ZT3 in VHS. In contrast, aerobic glycolysis was enhanced in VHS at ZT8 in pilo mice.

## DISCUSSION

The present work’s main findings are that oxidative metabolism and aerobic glycolysis are recruited differently in hippocampal regions involved in different cognitive functions. This recruitment is time regulated (ZT3 versus ZT8) in control mice, and it is altered in experimental epilepsy. VH displays a larger dynamic range for energy production than DH in both control and epilepsy conditions.

For this study, we chose an ex vivo approach because it allowed controlling the energy substrate composition and their delivery rate. It also allowed standardizing the electrical responses for all explored conditions. The various limitations of the experimental approach are discussed in SI Discussion section.

### Metabolic activity in the dorsal and ventral hippocampus of control mice at ZT3

At the beginning of network activation, both dorsal and ventral parts rely on oxidative metabolism before enhancing aerobic glycolysis as metabolic demand increases with activity. Both oxidative metabolism and aerobic glycolysis are larger in the ventral than in the dorsal part. Adding to the dissociation, the dorsal part relies longer on oxidative metabolism. The enhanced metabolic activity in the ventral part may appear surprising since the electrophysiological responses were similar in both dorsal and ventral parts. The local field potential mostly reflects the activation of synapses, while the metabolic measurements comprise both postsynaptic and presynaptic events as well as glial cell activity. A presynaptic mechanism may partly account for the dorsoventral difference. Although the probability of presynaptic release of both GABA and glutamate is greater in the ventral than in the dorsal hippocampus, the network response to Schaffer collateral activation remains the same (21). A greater neurotransmitter release probability should increase the metabolic load (e.g. for neurotransmitter recycling). Another contributing factor is the intrinsic excitability, which is greater in the ventral than in the dorsal hippocampus (11, 13, 21). This is consistent with the greater level of metabolic activity in ventral slices in the absence of stimulation. Besides, glial cells will be highly mobilized, particularly for K^+^ and neurotransmitter uptake (40). The VH contains 1/3 less astrocytes than the DH in rats (41). Therefore, the heavier functional load must be carried by a smaller quantity of astro-glial cells, which requires more intense metabolic activity. Beside ATP production, glucose metabolism is involved in other critical functions, including oxidative stress management, synthesis of neurotransmitters, neuromodulators, and structural components (3). Thus, the latter processes may also be regulated in a spatio-temporal manner.

Although glucose and oxygen consumption started simultaneously at the onset of synaptic stimulation in VHS, the onset of glucose consumption was delayed by 9 s compared to oxygen in the DHS. Alternative substrates include lactate, amino acids, and glycogen. Lactate can be ruled out because we would have measured a decrease in lactate concentration at the beginning of the stimulation, while the reverse was observed (Fig. 2C and E). Astrocytes can metabolize amino acids, especially glutamate as energy substrates (42). This scenario is unlikely as, at the beginning of the stimulation, their extracellular concentration should be low. Glycogen is a probable candidate: it is accumulated mainly in astrocytes (43), but it can also be found in smaller quantities in neurons (44). A larger astrocyte number in DH (41) may account for a preferential glycogen utilization in the dorsal region. The hippocampus is the most glycogen-rich and glycogen-consuming brain region (45). Therefore glucose needs to be utilized to replenish its store, in line with the continuous rise in glucose consumption after the end of the synaptic stimulation while oxygen consumption is declining, especially in the DHS. The reliance on glycogen in the DH may be linked to the functions performed in this region. Memory processes that are more specific to the DH may require glycogen. Some authors suggest the importance of glycogen derived lactate (46, 47), although lactate is rather ineffective to fuel synaptic transmission (48). Another hypothesis proposes glycogen utilization by astrocytes to spare glucose for energy-demanding neuronal functions (49).

Together, our results show a clear heterogeneity in energy metabolism between the ventral and dorsal parts of the hippocampus at ZT3. This heterogeneity may be linked to the fact that these two regions are implicated in different brain networks and functions (6–8).

### Circadian regulation of metabolic activity in control mice

Our results show enhanced oxidative glycolysis at ZT8 as compared to ZT3 in particular in VHS. Previous works indicate that both NAD(P)H dip and FAD^+^ peak reflect the same metabolic processes (39, 50). Yet, we found an increase in FAD^+^ and a decrease in NAD^+^ at ZT8 (Fig.8A, B). This discrepancy can be resolved by the changes in glucose consumption and lactate release, which were larger and smaller at ZT8 than at ZT3, respectively. Importantly, we often observed a decrease in lactate concentration at the beginning of the synaptic stimulation suggesting its consumption. Astrocytic lactate transport is not operational at the measured concentration of 0.1-0.3 mM while neuronal transport is (51). In neurons, lactate is converted to pyruvate by the lactate dehydrogenase (LDH), reducing NAD^+^ accounting for the observed decrease in NAD(P)H dip amplitude. Because the activity of the G3PS could be low at ZT8 (see SI Discussion) or because LDH activity may be coupled with malate-aspartate shuttle (52) this change in NADH does not affect the mitochondrial FAD^+^/FADH_2_ pool. In parallel, the rapidly available pyruvate will boost the Krebs cycle, increasing NADH. This would explain why NAD^+^ dip and FAD^+^ peak amplitudes go in opposite directions and argue for increased oxidative metabolism at ZT8. The oxidative shift may be also driven by the endogenous oscillation in redox state recently found in hippocampal slices (53).

### Metabolic activity in slices of dorsal and ventral hippocampus of epileptic mice

Our results show that the spatial and temporal dimensions are important to consider to understand how hippocampal cells produce energy in control animals. Investigating this issue in a pathological context is equally important as metabolic alterations may disrupt cell function, leading to cell death and participating in functional deficits, such as memory and learning. Epilepsy is a prototypical example of a pathological condition in which metabolic alterations have been identified (28). The differences between VH and DH were abolished at ZT3 in experimental epilepsy. Since we did not perform a systematic circadian analysis in control animals, we cannot rule out the possibility that such a phenomenon occurs at another ZT in control conditions. Said differently, what we observed at ZT3 in epilepsy would correspond to a phase shift of what happens at another ZT in controls.

We also noted a transient increase in glucose concentration at ZT3 after the stimulation onset. This increase cannot correspond to glucose release, because unlike endothelial cells, glial cells and neurons rapidly phosphorylate glucose to glucose-6-phosphate, thus trapping the substrate in the cell (1). The most likely possibility is a transient decrease in glucose uptake. The first step in glycolysis is glucose phosphorylation, which requires cytosolic ATP. At the onset of the synaptic stimulation, this ATP is also consumed by the Na^+^/K^+^ ATP-ase pump. In experimental epilepsy, ATP levels are 25% lower (54), and the slicing further decreases ATP levels by 40-60% (55), making the intracellular ATP concentration in hippocampal slices from pilo animals as low as 0.9-1.0 mM (see explanation in SI Discussion). Since the hexokinase has a lower affinity for ATP (K_m_=0.1-0.7mM, (56)) than the Na^+^/K^+^ ATP-ase (K_d_=0.1-0.2 μM, (57)), ATP will be preferentially used by the pump rather than by the hexokinase, resulting in decreased glucose metabolism. Increases in glucose concentration were more often observed in dorsal (7 from 9) than in ventral slices (3 from 9), which is consistent with the increased excitability of dorsal CA1 pyramidal cells in the pilocarpine model (58).

Together, the small amplitude of NAD(P)H and FAD^+^ signals, glucose, and lactate transients, as well as the initial positive deflection of glucose signal, suggest a deficient energy metabolism in both DH and VH in experimental epilepsy, in keeping with the hypometabolism of glucose found in the pilocarpine model of TLE (59–61). Importantly, the deficit was more pronounced in the VHS than in the DHS. This result is consistent with the fact that in this epilepsy model the neuronal loss, synaptic remodeling, and seizure initiation are more pronounced in the VH (62, 63)

At ZT8, when seizure probability is high, all metabolic signals revealed enhanced aerobic glycolysis as compared to ZT3, the increase being more prominent in the VH. Thus, the time of the day appears as a key parameter to take into account when looking at metabolism in epilepsy, particularly for diagnosis purposes in patients (64). In epileptic regions, glucose uptake is low during the interictal period (hypometabolism), but it increases during a seizure (65), presumably to fuel it. We have shown that a local change in aerobic glycolysis can occur over time without an associated change in network activity. We can speculate that metabolism waxes and wanes in a circadian manner, which may periodically drive networks close to seizure threshold (66).

In conclusion, our results demonstrate that both time and space are key parameters to consider for energy metabolism. Many brain regions develop in a dorsoventral manner (67), leading to heterogeneity in neuron and astrocyte spatial distribution and properties. The dorsoventral differences in energy metabolism may reflect such diversity as cells are integrated into distinct networks to perform different tasks. The time-dependent regulation of metabolism is the second important variable. Body functions are regulated by circadian clocks, which constantly change the molecular organization of cells and circuits. The circadian regulation of metabolism may just be part of this general principle, adapting metabolic needs to cell demands.

In epilepsy, the basic rules of a spatiotemporal pattern are still present, but they are considerably modified compared to the control situation. Whether such modifications are causally related to seizures and epilepsy comorbidities remain to be determined.

## ACKNOWLEDGEMENTS

Supported by A*MIDEX: 2IONXXID/REID/ID17HRU208 and ANR-17-GRF2-0001-03.

## AUTHOR CONTRIBUTIONS

Conceptualization, C.B.; Methodology, A.G. and A.I.I.; Investigation, G.E.B., C.R., A.G. and A.I.I.; Formal Analysis, Visualization and Writing –Original Draft, A.I.I. and A.G.; Validation and Writing –Review & Editing, C.B. and A.I.I.; Project Administration and Funding Acquisition, C.B.; Supervision, C.B and A.I.I.

