## Supplementary information for "Spatio-temporal heterogeneity in hippocampal metabolism in control and epilepsy conditions"

**This PDF file includes:**

Detailed description of Methods
SI Discussion
Limitation of the approach used
Local cerebral glucose utilization in dorsal and ventral hippocampus *in vivo*
Estimation of ATP concentration in brain slices.
Activity of glycerol-3-phosphate shuttle in hippocampal slices.
Figures S1 to S4
Tables S1 to S4
SI References

**Supplementary Information Text**

### METHODS.

*Tissue slice preparation*

All animals were kept in colonies to prevent single housing-induced stress (1) . To avoid the heterogeneity in slice properties that might be induced by a seizure, animals' behavior was observed 30 minutes before euthanasia to be free of seizure. Adult (2-4 months) FVB male mice were anesthetized with isoflurane before decapitation at Zeitgeber 3 and 8 (i.e. 10:00 and 15:00, respectively). The brain was rapidly removed (< 30s) and placed in ice-cold artificial cerebrospinal fluid (ACSF). The ACSF solution consisted of (in mM): NaCl 126, KCl 3.5, $NaH_2PO_4$ 1.25, $NaHCO_3$ 25, $CaCl_2$ 2, $MgCl_2$ 1.30, and glucose 10, pH 7.4. ACSF was saturated in 95% $O_2$ / 5% $CO_2$ gas mixture. After cerebellum removal, the two hemispheres were separated. One hemisphere, left or right in alternation, was used to prepare dorsal slices, whereas the other hemisphere was used to prepare ventral slices as described previously (2). 350 µm thick slices were obtained with a Leica VT 1200s (Leica Microsystem, Germany). During cutting, slices were submerged in an ice-cold (< 6°C) solution consisting of (in mM): K-gluconate 140, HEPES 10, Na-gluconate 15, EGTA 0.2, NaCl 4, pH adjusted to 7.2 with KOH. Slices were immediately transferred to a dual-side perfusion holding chamber with constantly circulating ACSF and allowed to recover for 2 h at room temperature (22°C-24°C).

*Synaptic stimulation and field potential recordings.*

After recovery, slices were transferred to a dual perfusion recording chamber continuously superfused (6 ml/min) with ACSF contained 5mM of glucose (see explanation below), warmed to 30°C. Schaffer collateral/commissures were stimulated with a bipolar metal electrode (Fig.S1, 'SE') connected to a DS2A isolated stimulator (Digitimer Ltd, UK). The electrode tip was placed in *stratum radiatum* between the CA3 and CA2 areas. The extracellular local field potential (LFP) was recorded with a glass microelectrode filled with ASCF, placed in *stratum pyramidale* of the CA1 area (Fig.S1, 'RE'). The signal was amplified

1000 times with an EXT-02F amplifier (NPI Electronic, Germany) operating with a low pass 3kHz filter (DC mode).
We first increased the stimulation intensity (applied at a frequency of 0.15 Hz, using a fixed duration of 200 µs) until reaching the maximum amplitude of the LFP. The current intensity was then adjusted to produce a LFP response at 50-60% of the maximal amplitude. We used this current intensity during trains of stimulations applied at 10 Hz during 10 or 30 s. We tested different train frequencies and found that 10 Hz was optimal because i) it did not damage the slice tissue, even after several (10 trains) stimulation sessions, since it produced identical responses when applied at least 10 times every 5 minutes, ii) it generated LFP responses on each pulse of the train (Fig.S2 A), iii) it did not trigger after-discharges, iv) it induced transient change in all metabolic parameters with an amplitude significantly higher (>5 times) than a standard deviation of the baseline (30s before stimulation).  Besides, the 10 Hz frequency is within the theta oscillation range and the seizure discharge range. The 10 s and 30 s durations are also within the range of theta episodes and seizures.
The stimulation intensity necessary to evoke a 50% maximum LFP response was equally variable between 140-300 µA in dorsal and ventral slices (Fig.S2 Ba). To assess the response to the stimulation train, we calculated the integral of each LFP response in the train (Fig.S2 Bb) as well as the cumulative integral of all the responses (Fig.S2 Bc), following a previously established procedure (3). Figs.S2Bb and S2Bc show no difference in evoked electrophysiological responses in ventral and dorsal slices. This allowed standardizing the network response in dorsal and ventral slices, even if their intrinsic excitability may be different (see SI Discussion).

*Oxygen, glucose, and lactate measurements.*
We used a Clark-style oxygen microelectrode (tip diameter 10 µm; Unisense Ltd, Denmark) to measure slice tissue $pO_2$. We set the electrode polarization at -0.8 mV and measured the generated current with a PA2000 potentiostat (Unisense Ltd, Denmark). Tissue glucose and lactate concentrations were measured with enzymatic microelectrodes (sensing tip diameter 25 µm, length 0.5 mm; Sarissa Biomedical, Coventry, UK) polarized at 0.5V and driven with the free radical

analyzer TBR4100 (WPI, USA). The sensing element of the enzymatic electrode was a platinum/iridium wire coated by a layer of analyte-specific enzyme and encapsulated in an analyte selective membrane. The enzyme catalyzes the oxidation of the substrate, producing $H_2O_2$, which reduction on the surface of platinum/iridium wire provides electrons for the current driven by the polarization potential. Thus, all enzymatic electrodes, independently of their analyte, produced and sensed $H_2O_2$. Since their selective membrane is permeable to $H_2O_2$ some of the produced $H_2O_2$ unavoidably diffuses to the tissue or another enzymatic electrode located nearby. To minimize possible toxic effects of $H_2O_2$ when using the glucose sensor, we used ACSF containing glucose at 5 mM instead of 10 mM. With 5mM glucose and a high perfusion rate (6ml/min, and 1 ml chamber volume), the ambient lactate concentration in the slice was 50-250 µM, i.e. nearly two orders of magnitude lower than glucose. At this concentration, the quantity of $H_2O_2$ produced by the lactate sensor and diffusing in the tissue is very low (i.e. the toxicity is negligible). In each experiment, we monitored slice viability looking at LFP responses' stability to single-pulse stimulation. Slices were changed if the response amplitude changed by more than 20%.
To avoid an interaction between enzymatic glucose and lactate electrodes, glucose and lactate measurements were done in separate sets of experiments. However, oxygen measurements could be done simultaneously with glucose or lactate ones as the corresponding electrodes have a different design.
Glucose or lactate measurements could not be performed simultaneously with NAD(P)H or $FAD^+$ because the electrodes' sensitive site contains flavin adenine dinucleotide (4), the enzymatic sensors are sensitive to the excitation light applied for NAD(P)H or $FAD^+$ imaging.

All electrical signals (LFP, glucose-, lactate-, $O_2$-generated currents) were digitized using InstruTECH LIH 8+8 acquisition interface (HEKA Electronics, Germany) and stored on the computer hard drive. Stimulation and recording controls were driven with the Patchmaster software (HEKA Electronics, Germany). A second acquisition system, MiniDigi1B interface card and Axoscop software (Molecular Devices, CA), was used to record all signals at a 1 kHz time resolution during the whole duration of the experiment. These recordings were used to estimate the change of sensor sensitivity as a decrease of the baseline current generated by the sensors between synaptic stimulations. The oxygen sensor remained stable for several days, whereas enzymatic electrodes lost 20-30% of their sensitivity at the end of each experiment. Continuous recordings were used to estimate the drift in sensitivity to correct lactate and glucose measurements with an exponential function. Between experiments, the enzymatic electrodes were cycled in analyte free ACSF from -500 mV to + 500 mV and back at a rate of 100 mV/s for 3 cycles. They were then polarized to +500 mV during 30 min and calibrated at 5 points in ACSF warmed to 30°C. Glucose and lactate sensors were calibrated before and after each recording session.

*NAD(P)H and $FAD^{\pm}$ fluorescence imaging.*

NADPH and NADH have similar optical properties. Therefore, it is expected that NADPH may contribute to the total autofluorescence signal. Because the cellular $NADP^{+}$/ NADPH pool is about one order of magnitude lower than the NAD/NADH pool, we assume that short-term variations of experimentally detected fluorescent changes responses account in the present study for variations in NADH (5) . In the present study, we use the term NAD(P)H for the fluorescent signal.

Slices were epi-illuminated with monochromatic light (pE-2 illuminator, CoolLed, UK), using 365 nm for NAD(P)H and 440nm for $FAD^{+}$. The power of illumination was adjusted to excite the whole thickness of the slice (30-40% for NAD(P)H and 40-50% for $FAD^{+}$ of the maximum power of the LED). The emitted light was filtered by 420 nm or 500 nm long-pass filter (Omega Optical, Brattleboro, VT) and imaged through a Nikon upright microscope (FN1, Eclipse) with 4x/0.10 Nikon Plan objective using a 16-bit Pixelfly CCD camera (PCO AG, Germany). Because of the low level of fluorescence emission, NAD(P)H and $FAD^{+}$ images were acquired every 600-800 ms as 4x4 binned images (effective spatial resolution of 348x260 pixels). The exposure time was adjusted to obtain baseline fluorescence intensity at 50% of the camera dynamic range. Fluorescence intensity changes were measured in *stratum radiatum* near the site of LFP recordings in 3 regions of interest ~1mm distant from the stimulation electrode tip using ImageJ software (NIH, USA). Data were expressed as the percentage changes in fluorescence over a baseline [(ΔF(t)/F(t)) • 100%] were F(t) is an exponential function of time approximating the base line fluorescence decay estimated using the recording of 1 min base line fluorescence preceding the stimulation (for more details see (6)). Signal analysis was performed using IgorPro software (WaveMetrics, Inc, OR, USA).

*Experimental model of epilepsy*

Adult FVB mice were injected with methylscopolamine (1 mg/kg i.p.) 30 min before the pilocarpine injections. Pilocarpine was repeatedly injected (100 mg/kg i.p.), every twenty minutes until status epilepticus (SE) was observed. After 90 min of SE, we injected diazepam (10 mg/kg i.p.) to stop SE. All mice then received 0.5 ml NaCl (0.9%) subcutaneously and again in the evening. During the following days, if required, mice were fed with a syringe.

At chronic stage, pilo mice weighed less than control or sham (27.1±2.8 g n=26, 33.5±1.8 g n=9, 31.5±1.7 g n=14, respectively p<0.0001 ANOVA). Still, they did not present any signs of severe or moderate suffering (the averaged MGS score estimated according to (7) was 0.8 ±0.2 AUs). All mice which experienced status epilepticus (n=21) survived. Pilo mice were not instrumented with EEG telemetry to keep perturbations to a minimum. However, in this model, telemetry measurements performed in another batch of animals show that all pilo mice experiencing status epilepticus develop spontaneous seizures with a low incidence in the morning (2.5-4% between ZT1-3), and a high probability in the afternoon, especially at ZT8 (11%) (8)

*Statistical analysis and signal processing*

To evaluate the difference between metabolic parameter values measured in dorsal and ventral slices, we used in parallel two statistical approaches: the non-parametric significance tests and estimation statistic of bootstrap samples. The first approach calculates the probability that the difference would be observed if it does not exist. Using Shapiro-Francia normality test, we verified if the experimental data set could be considered normally distributed. If yes, we

applied two tails unpaired or paired t-test. Otherwise, Mann-Whitney (M-W) or Wilcoxon (Wilc) tests were used to compare unpaired or paired data. Although commonly used, this approach is dichotomous, so it allows only to answer if the difference exists or not without estimating how large the difference is. The second approach consisted of calculating the 95% confidence interval (95%CI) of the mean difference by performing bootstrap resampling (9). It resamples (with replacement) a single set of observations and calculates the difference between the values of a statistical parameter (the mean in this study) for compared groups (VHS-DHS or ZT8-ZT3). After repeating this procedure several times (5000 in our case), the distribution of differences is computed. The mean of this distribution is an estimator of the size of the difference, while its 95%CI is a measure of the precision and confidence of this difference. In the text, we reported the difference size as: unpaired or paired *mean difference value [lower bound, upper bound of 95%CI]*. In some cases, when the absolute values were of interest, we reported the corresponding parameter name and its mean value with 95% confidence interval obtained using the bootstrap procedure.
This type of analysis was performed using the estimation statistics website www.estimationstats.com, the accuracy of the results was verified with an R language script. In both cases the replication number was 5000, and confidence interval was bias-corrected. For an illustration of data analysis, we used estimation plot (10) which shows experimental data set as circles plotted on left hand side Y-axis (black color) and the distribution of estimation parameter (usually ventral -dorsal mean difference) of the bootstrap samples is plotted as gray bell shape area on a floating right hand side (gray color) Y-axis. The mean value of this distribution (black closed circle) as well as its 95% bias corrected confidence interval (black error bar) are also plotted on right hand. The results of significance tests, *p* values, are reported on figures. The asterisk code: *- $0.01<p<0.05$, ** - $0.001<p<0.01$, *** $p<0.001$. We used a two-way ANOVA to assess the variability of optical imaging parameters with time.

### DISCUSSION

#### *Limitations of the study*

We used stimulation trains to assess energy metabolism. To remain close to physiologically relevant conditions, we used a 10 Hz frequency, which is within the range of theta oscillations (4-12 Hz), a brain rhythm associated with numerous cognitive and behavioral functions (11) and lengths (10 s and 30 s) that are compatible with the duration of locomotion and REM sleep episodes, for example. The 10 Hz frequency is also within the 0.5-40 Hz band of oscillations found during seizures, lasting between 10 and 30 s. Although artificial, the trains used in the present study are consistent with the synaptic activity CA1 pyramidal cells would experience in physiological and pathological conditions. Electrical stimulation allowed us to normalize the electrophysiological responses, enabling a direct comparison between the dorsal vs /ventral and ZT3 vs ZT8 conditions in control animals. However, we could not directly compare the evoked responses between control and epilepsy conditions since the electrophysiological responses were systematically smaller in pilo than in control animals, probably due to the cell loss that characterizes this model (12). Pharmacological approaches can be used to evoke activity in slices, including the perfusion of carbachol or kainic acid to trigger theta and gamma oscillations (13, 14), respectively, but oscillations are very variable from one slice to another, preventing normalization across conditions.

Brain cells must have access to nutrients (glucose, oxygen) and the possibility to eliminate waste products to maintain a healthy state. In the *ex vivo* brain preparation, this goal is achieved thanks to the high-rate perfusion of the slice with ACSF on both sides of the slice, which prevents the creation of the zone of hypoxia and hypoglycemia in the bottom part of the slice, especially during the strong metabolic activity associated with the neuronal network stimulation (15) . We showed previously that in these experimental conditions, the amplitudes of the NAD(P)H dip and the $FAD^+$ peak are associated with the activity of oxidative metabolism because they are directly related to the oxygen consumption, whilst the overshoot and undershoot components are associated with the activity of cytosolic glycolysis because they are abolished if cytosolic glycolysis is blocked when substituting glucose for pyruvate (3, 6, 16) see also (17)). When a lower perfusion rate on one side only is used, the center and bottom slice are in hypoxic and hypoglycemic conditions, reducing oxidative phosphorylation leading to an increase in mitochondrial NADH and $FADH_2$. Consequently, the $FAD^+$ and NAD(P)H signals are mainly of mitochondrial origin (18), so they are not comparable with those presented in this work.

We chose an *ex vivo* approach because it is not possible to standardize the stimulation in freely moving animals and because oxygen and glucose supply cannot be clamped *in vivo* due to the variable hemodynamic. Also, the absorption spectra of oxygenated and deoxygenated blood are different, so the transition from one to the other can affect the NAD(P)H and $FAD^+$ signals (19). Besides, invasive equipment would induce astrogliosis and neuroinflammation, which would alter the metabolic state of the brain tissue.

Finally, the methods do not offer enough resolution to distinguish between neuronal and glial cell effects. The oxygen electrode can measure metabolic changes in a small tissue volume comprising a few cells (sphere of 30 µm in diameter). All others techniques used here integrated metabolic signals over a much larger volume, which was equal to the thickness of the slice. This difference in spatial scale is probably responsible for some discrepancies observed between the oxygen detection results and all the other methods. However, the techniques used here allowed us to explore the same metabolic events from the intracellular (NAD(P)H and $FAD^+$ imaging) and extracellular sides. By matching the intra- and extra-cellular signals, we obtained important conclusions about the metabolic specificity associated with a difference in space (dorsal vs. ventral), time (ZT3 vs. ZT8), and functional state (healthy vs. epileptic brain). The scheme that we propose does not distinguish between neurons and glial cells; thus, we summarize our results on a "neuro-glial" cell (Fig.S3).

We used dorsal and ventral slices, i.e. two gross spatial locations along the hippocampus's septo-temporal axis. While some coronal slices can give access to both septal and temporal regions simultaneously, the Schaffer collaterals' wrong orientation prevents using the same stimulating protocol. Future studies are needed to determine whether there is a smooth gradient of metabolic properties along the longitudinal axis, as demonstrated for membrane oscillations in entorhinal stellate cells (20), or whether specific domains exist as demonstrated for gene expression along the longitudinal axis of the hippocampus (21). Also, we performed our measurements in distal CA1, far from the stimulating electrodes. Since there is morpho-functional gradient along the transverse axis (22) (from proximal CA1 close to the CA2 to distal CA1 close to the Subiculum), we cannot rule out that the properties of metabolism also vary along this axis. Likewise, these properties may differ in the other subfields (CA2, CA3, dentate gyrus) of the hippocampus.

We used only two time points – ZT3 and ZT8 – because they correspond to a low and high seizure probability in pilo mice, respectively. Yet, even with two time points, we could already evidence differences in control animals, thus providing the proof-of-concept of a circadian regulation of metabolism in the CA1 region of the hippocampus. Further studies are needed to assess metabolic function hour per hour, particularly during the night (active) phase.

We did not instrument pilo animals to monitor seizures. Hence, spontaneous seizures occurring prior to measurement may have altered energy metabolism. We decided not to instrument mice to remove two confounding factors: the unknown effects of invasive surgery and anesthesia and the stress induced by single cage housing required for instrumented animals. Single cage housing produces a very severe epilepsy phenotype, while group housing decreases seizure frequency by at least one order of magnitude (1). Although we cannot rule seizure-induced metabolic alterations, we could still evidence spatio-temporal differences in pilo mice. Obviously, we cannot claim a homology between an animal model and human epilepsy. We were only interested in the broad context of a neuronal network generating spontaneous seizures.

Despite these limitations, our results reveal clear dorso-ventral, time, and condition (control/epilepsy) dependent energy metabolism changes.

*Local cerebral glucose utilization in dorsal and ventral hippocampi in vivo.*

Studies performed *ex vivo* must be interpreted with care as the environment of cells differs from the one in the intact organism. In this context, it is particularly important to compare our results to those obtained *in vivo.* We found few studies where the local cerebral glucose utilization rate(LCGU) was evaluated in DH and VH of conscious rats using [$^{14}$C] 2-deoxy-D-glucose autoradiography (23-28) . The values are summarized in Table S4. In five of the 6 reports, the LCGU was 17 ± 3% (mean±sem) higher in the ventral hippocampus than in the dorsal hippocampus all structures combined, and 28 ± 3% (n=4) higher in ventral than dorsal CA1. One work reported identical LCGU values (28) , but this study was performed on young animals (P21). Interestingly, we found a similar VH – DH dissociation *ex vivo*: the steady-state glucose utilization in the CA1 was also higher by 23±5% in ventral than dorsal slices. Although absolute metabolic values are likely very different between *in vivo* and *ex vivo* conditions, ratios may be maintained.

*Estimation of ATP concentration in brain slices.*

Our estimation is based on the data published by Sonnewald group (29). Using $^{1}$H nuclear magnetic resonance spectroscopy, the authors measured ATP+ADP concentration in rat brain extracts of the hippocampal formation, cerebral cortex, and cerebellum from chronically epileptic rats (pilo model) studied during their interictal phase and compared the values with age- and weight-matched controls. They found that, in the hippocampus, ATP+ADP levels were 25% lower in epilepsy than in control (2.07 ± 0.46 µM/g of tissue vs. 1.55 ± 0.29µM/g of tissue, respectively). According to zur Nedden and colleagues (30), the slicing procedure decreases the total adenine nucleotide pool (TAN) and ATP to ADP ratio. These parameters recover partially during incubation, and 1-3 hours after cutting (time corresponding to our experiments) the TAN in slices represents 40-44% of TAN in the intact hippocampus, while ATP to ADP ratio increases from 8 to 12. By taking into account these values as well as the volume of the slice ($8.75x10^{-3}$ ml) and its dried weight (14 mg), we estimated that the upper limits of intracellular ATP concentration to be 1.17-1.34 mM and 0.90-1.02 mM in control and epilepsy conditions, respectively. Since a fraction of this ATP may be located in secretory vesicles, the ATP amount available for cytosolic enzymes, such as hexokinase, should be lower.

*Activity of glycerol-3-phosphate shuttle in hippocampal slices.*
The results of our imaging experiments suggest the coupling between cytosolic and mitochondrial redox state. As is shown on Fig.1 and Fig.5 the increase in stimulation duration produced symmetrical changes in NAD(P)H and $FAD^+$ wave shapes. This coupling may be carried out by the activity of the glycerol-3-phosphate shuttle (G3PS). The G3PS is formed by two glycerol-3-phosphate dehydrogenases, namely cytosolic cGPDH that is soluble and NADH-dependent (E.C.1.1.1.8), and by mitochondrial, membrane-bound, FAD-dependent mGPDH (E.C. 1.1.5.3). They catalyze the conversion of dihydroxyacetone phosphate to glycerol-3-phosphate and vice versa, coupled with oxidation of cytosolic NADH and reduction of mitochondrial $FAD^+$. The main metabolic role of this shuttle is the reoxidation of cytosolic NADH produced by glycolysis. This enables a sustained cytosolic ATP production rate without accumulating an intermediate by-product such as lactic acid. This mechanism also leads to the transport of reducing equivalents (hydrogen) from the cytosol into mitochondria, where they can be utilized for aerobic mitochondrial ATP production (31) . However, there are contradictory data about the role of G3PS in brain metabolism (reviewed in (32). Indeed, G3PS was not considered of general importance because the functional activity of the shuttle requires equimolar proportions of both components  - cGPDH and cGPDH. In most tissues, including the brain, the content of mGPDH is relatively low compared to cGPDH (31). Therefore, in the brain, G3PS appears to be only secondary to the malate/aspartate-shuttle. However, it was shown that the brain can metabolize exogenous glycerol *in vivo*;

a process that requires the activity of mGPDH (33). Moreover, in the same study using histochemical staining, the authors demonstrated the expression of mGDPH in areas of high synaptic density, e.g. in *stratum oriens, stratum radiatum*, and molecular layer. They also provided indirect evidence that glycerol is metabolized more by neurons, especially GABAergic ones, than by astrocytes. Also, there is rapid biosynthesis of glycerol-3-phosphate in response to electrical stimulation of Schaffer collateral (34) – a stimulation protocol similar to that used in the present work. By analyzing metabolic fluxes, the authors concluded that G3PS is not active in baseline conditions but is boosted after synaptic stimulation, especially in neuronal cells. Thus, there is strong evidence for the activity of G3PS in the hippocampus that can explain the observed coupling between cytosolic NADH and mitochondrial $FADH_2$ in control conditions.

Such a coupling was not observed in slices from epileptic mice made at ZT3 (main text Fig.3), and control slices made et ZT8 (main text Fig.8). Since hippocampal genes are regulated in a circadian manner and since this regulation is modified by epilepsy (8), we checked in the CircadiOmics database (35, 36) whether the activity of genes of proteins involved in G3PS were regulated in a circadian manner in the mouse hippocampus. We found that the expression of Gpd1, the gene responsible for encoding cGPDH, is lower at ZT8 than at ZT3 in the ventral hippocampus of control mice (experiments were not performed in the dorsal hippocampus). In contrast, in epileptic mice, this protein is more expressed at ZT8 than at ZT3 (Fig. S4). This finding is in agreement with our results obtained using NADH and $FAD^+$ imaging. Together, our data and already published results suggest that G3PS is operational in the mouse hippocampus during synaptic activation and that its activity is involved in the redox coupling between the cytosol and mytochondria, in a circadian dependent manner.

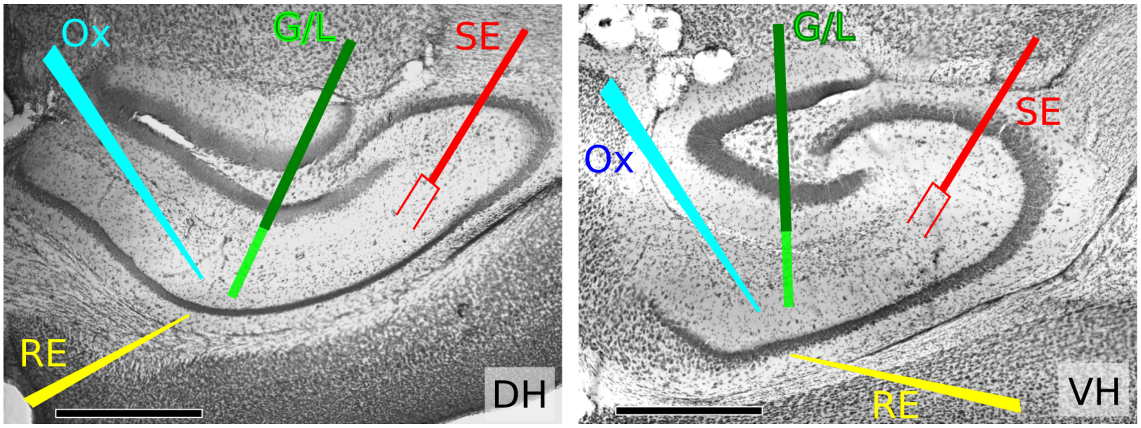

***Fig.S1 Photomicrograph of Nissl stained 60µm thick hippocampal slices cut from dorsal (DH) and ventral (VH) regions.*** *The overlays indicate the positions of recording electrode (RE) stimulating electrode (SE), oxygen sensitive electrode (Ox) and glucose or lactate enzymatic sensor (G/L). Black bars in bottom left corners are 500µm scale bars.*

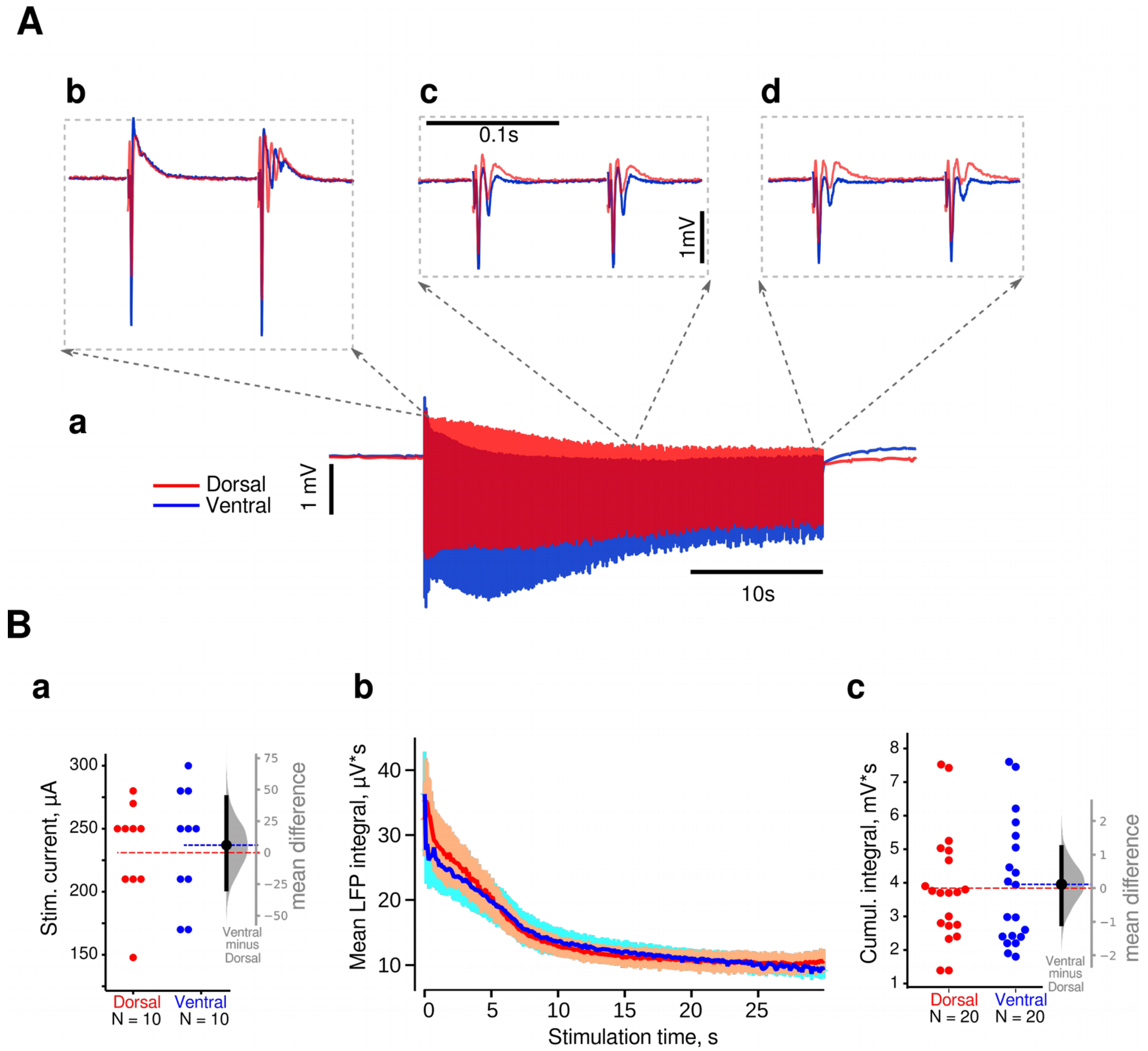


***Fig. S2. The stimulation of Schaffer collaterals (10 Hz during 30 s) induced similar LFP responses in slices from dorsal (red traces) and ventral (blue traces) hippocampal regions.***

***A. a.*** *Example of an entire LFP response evoked in dorsal and ventral slices. LFP responses were largest at the beginning of stimulation (**b**), decreased with time (**c**) but were still present at the end of the stimulation (**d**).* ***B.*** *Statistical summary of stimulation parameters and LFP responses to a stimulation train from 10 experiments (10 dorsal and 10 ventral slices, 2 stimulations per slice).* ***a:*** *the mean difference between the intensities of the current used to produce 50-60% maximum responses in ventral and dorsal slices was 6[-29, 44] µA, demonstrating similar properties regarding the stimulation intensity. b: Evolution of the mean integral for each LFP response in the train. The color lines represent the mean values, and the light color shading indicates SEM.* ***c:*** *Cumulative integrals of LFP responses (trains were given twice in slices) showed similar values in DH and VH. The mean difference between ventral and dorsal slices was 0.11[-1.07, 1.22]mV*s.*

*On **Ba** and **Bc** The mean difference between parameter values is shown as a Gardner-Altman estimation plot (see Statistical analysis section of Materials and Methods chapter). The mean difference distribution is plotted on floating gray axes on the right, as a bootstrap sampling distribution (light gray bell shape area). The mean difference is depicted as a black dot; the 95% confidence interval is indicated by the ends of the vertical error bar. Red and blue dashed lines correspond to mean values for dorsal and ventral slices. The 0 point of the mean difference is aligned to the mean value measured in dorsal slices.*

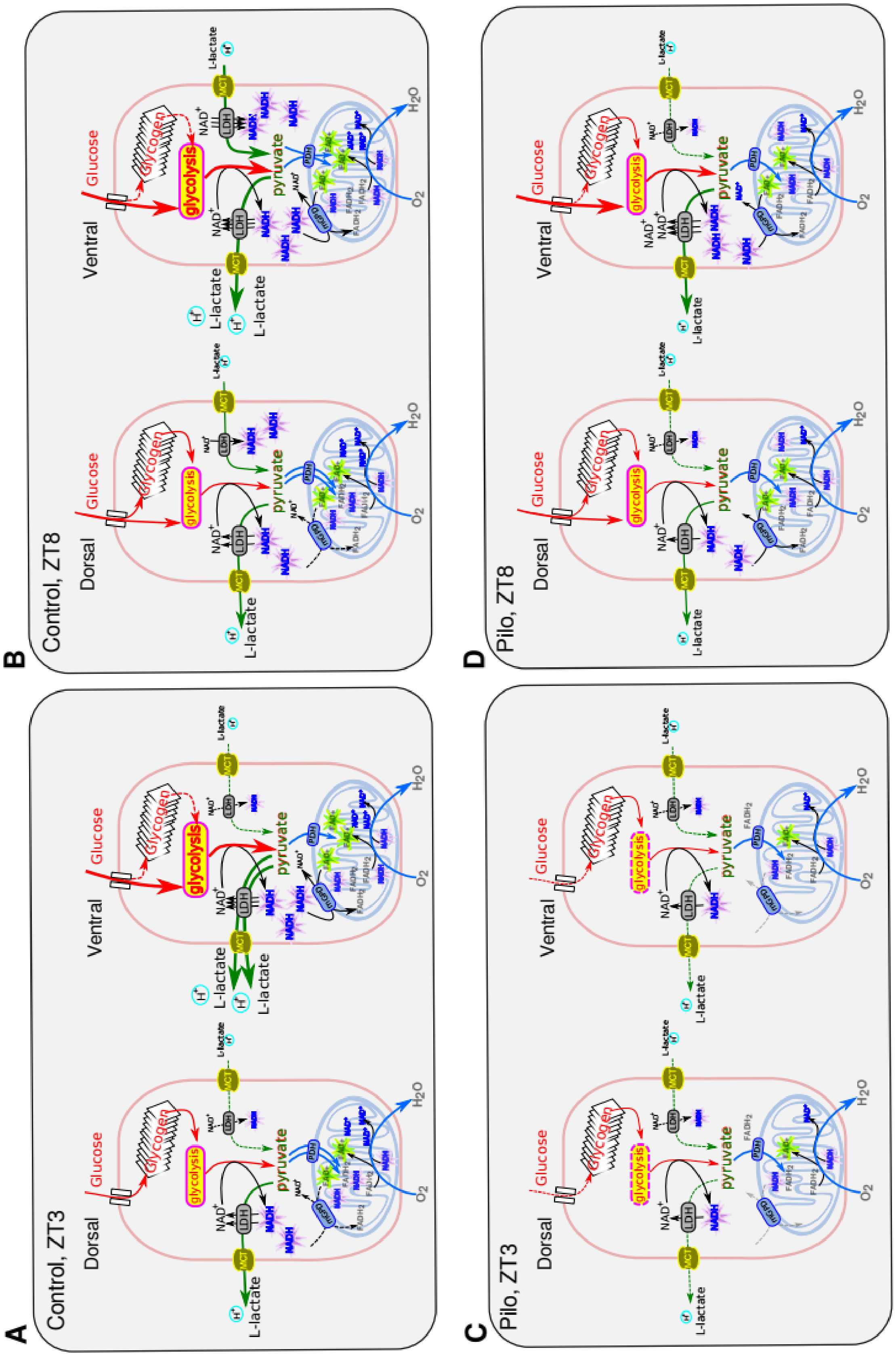
A
Control, ZT3
B
Control, ZT8
C
Pilo, ZT3
D
Pilo, ZT8
Dorsal
Ventral
Glucose
Glycogen
glycolysis
pyruvate
L-lactate
LDH
MCT
PDH
NAD+
NADH
FADH2
O2
H2O

***Fig. S3 Cartoon illustrating regional (dorsal/ventral) and temporal (ZT3/ZT8) differences in energy metabolism in normal (Control) and epileptic (Pilo) conditions.***

***A.*** *In control mice, the activity of ventral hippocampal cells (neurons/astrocytes) is fueled by aerobic glycolysis, which consumes high glucose levels and produces high levels of NADH. Cytosolic NADH activates the lactate dehydrogenase (LDH)-driven conversion of pyruvate to lactate and the mitochondrial glycerol-3-phosphate shuttle (mGPD). Both pathways regenerate cytosolic* $NAD^+$ *required for glycolysis. In parallel, mGPD reduces mitochondrial* $FAD^+$*, increasing the amount of* $FADH_2$ *available for ATP production by oxidative phosphorylation. In dorsal hippocampal cells, glucose consumption is delayed and long-lasting, suggesting the utilization and replenishment of glycogen stores. The lactate production is lower, indicating that a large fraction of pyruvate is going to mitochondria via the pyruvate dehydrogenase complex (PDH). The overall metabolic activity in dorsal cell is lower than in ventral.*

***B.*** *At ZT8, the metabolic activity is increased, in particular in ventral cells. Not only glucose but also lactate are consumed at the onset of synaptic stimulation. The rise in cytosolic NADH resulting from lactate to pyruvate is oxidized by the malate-aspartate shuttle (not represented) that increases the mitochondrial NADH but not* $FADH_2$*.*

***C.*** *Energy metabolism activity is decreased in epilepsy, especially in ventral cells. The regional difference disappears at ZT3. Due to the low cytosolic ATP content, the hexokinase activity that reduces glucose utilization is low. Therefore, in both regions, the glycolysis is fueled by glycogen, which is abundant (37) but has an abnormal structure (38) that prevents its rapid breakdown. Due to the low glycolysis rate, there is no coupling between cytosolic NADH and mitochondrial* $FADH_2$*. The energy is produced mainly in mitochondria.*

***D.*** *At ZT8 the regional difference reappears. In both ventral and dorsal parts, aerobic glycolysis is enhanced, but in the ventral part glucose consumption and lactate release are larger than in the dorsal part.*

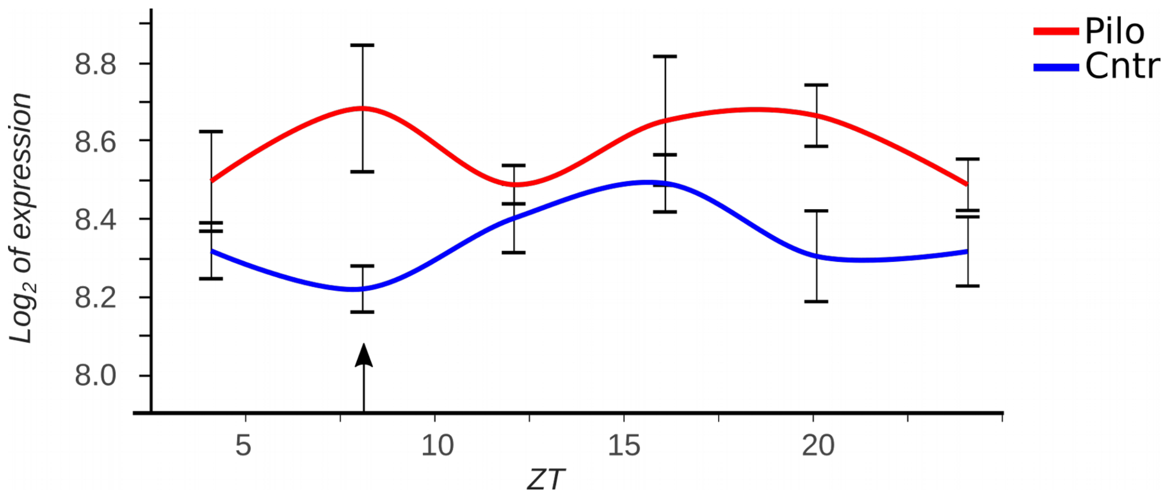


***Fig. S4 Time profile of the Gpd1 gene transcript expression in ventral hippocampus of control (blue) and Pilo (red) mice.*** *ZT-zeitgeber time in hours. Note at ZT8 (arrow) the transcript expression is minimal in control but maximal in pilo animals. Data on open access: http://circadiomics.igb.uci.edu/*

**Table S1.** Mean difference in amplitude of $O_2$, glucose, and lactate concentration changes induced by 10 s and 30 s stimulation trains in control slices.

In both slice types, the longer stimulation increased glucose and oxygen consumption similarly (see column Ventral vs Dorsal mean difference of difference), while the gain in lactate release was significantly higher in ventral slices.

| Parameter | Dorsal<br>30s vs 10s<br>Mean diff. [95%CI]<br>*p value* of Wilcoxon paired test | Ventral<br>30s vs 10s<br>Mean diff. [95%CI]<br>*p* value of Wilcoxon paired test | Ventral vs Dorsal<br>Mean diff of diff.<br>[95%CI] |
|---|---|---|---|
| Oxygen, µM, n=12 | 47 [36, 58]<br>*p=0.02* | 48 [31, 73]<br>p=0.008 | 0 [-20, 28] |
| Glucose, µM, n=7 | 56 [33, 104]<br>*p=0.04* | 87 [50, 123]<br>*p=0.007* | 31 [-21, 76] |
| Lactate, µM, n=5 | 33 [17, 54]<br>*p=0.03* | 81 [50, 118]<br>*p=0.005* | 49 [10, 90] * |

**Table S2.** Comparison of NAD(P)H and FAD+ transients recorded in slices of dorsal and ventral hippocampi of pilo mice at ZT3 using estimation statistic and Mann-Whitney U test

| NAD(P)H | | | $FAD^+$ | | |
|---|---|---|---|---|---|
| **Dorsal vs Ventral** | | | **Dorsal vs Ventral** | | |
| Parameter | Mean dif [95%C.I.] | M-W test, p | Parameter | Mean dif [95%C.I.] | M-W test, p |
| Dip amplitude | -0.1[-0.4, 0.2]% | 0.38 | Peak amplitude | -0.2[-0.6, 0.2]% | 0.49 |
| Overshoot amp.<br>10s<br>30s | <br>0.3 [-0.5, 1.4]%<br>0.8[-0.1, 1.6]% | <br>0.65<br>0.13 | Undersht amp.<br>10s<br>30s | <br>1.7 [-0.5, 3.7]%<br>1.4 [-1.9, 4.4]% | <br>0.13<br>0.41 |
| **10s vs 30s** | | | **10s vs 30s** | | |
| Parameter | Mean dif [95%C.I.] | Wilc. paired, p | Parameter | Mean dif [95%C.I.] | Wilc. paired, p |
| Overshoot onset<br>dorsal<br>ventral | <br>12 [8, 16]s<br>9 [2, 16]s | <br>0.02<br>0.03 | Undersht onset<br>dorsal<br>ventral | <br>2 [1, 5]s<br>4 [1, 7]s | <br>0.03<br>0.03 |
| Overshoot amp.<br>dorsal<br>ventral | <br>-0.1 [-0.6, 0.4]%<br>0.4 [-0.7, 1.5]% | 0.5<br>0.4 | Undersht amp.<br>dorsal<br>ventral | <br>-1.9 [-3.1, -1.0]%<br>-2.2 [-3.5, -1.0]% | <br>0.02<br>0.04 |

**Table S3.** Difference between metabolic parameters of slices prepared at ZT3 and ZT8 from pilo animals.

| | ZT8 vs ZT3<br>Mean diff. [95%C.I]<br>Mann-Whitney test *p value* | |
|---|---|---|
| Parameters | Dorsal | Ventral |
| Oxygen concentration change of amplitude | -5[-25, 17] µM<br>*p=0.8* | -27[-52, 1] µM<br>*p=0.2* |
| Glucose concentration change of amplitude | 0.1 [0.03, 0.28] mM *<br>*p=0.011* | 0.28 [0.14, 0.45] mM *<br>*p=0.006* |
| Glucose consumption onset | -5 [-7, -2] s *<br>*p=0.02* | -6 [-10, -3] s *<br>*p=0.06* |
| Lactate concentration change of amplitude (release) | 24 [-16, 58] µM<br>*p=0.3* | 58 [23, 89] µM *<br>*p=0.02* |

**Table S4.** Difference between metabolic parameters of slices prepared at ZT3 and ZT8 from control animals

| | ZT8 vs ZT3<br>Mean diff. [95%C.I]<br>Mann-Whitney test *p value* | |
|---|---|---|
| Parameters | Dorsal | Ventral |
| Glucose concentration change of amplitude | 0.11 [-0.12, 0.49]mM<br>p=0.08 | 0.5 [0.15, 0.86]mM*<br>p=0.17 |
| Glucose base line concentration | 1.4[1.0, 1.9]mM*<br>p=0.0002 | 1.32[0.6, 2.2]mM*<br>p=0.04 |
| Glucose consumption onset | 2[-3, 8]s<br>p=0.71 | 1[-1, 2]s<br>p=0.28 |
| Oxygen concentration change of amplitude | -44[-70, -20]µM*<br>p=0.007** | -18[-50, 25]µM<br>p=0.5 |
| Oxygen base line concentration | 72[11, 110]µM*<br>p=0.0006*** | 50 [-1, 84]<br>p= 0.036* |
| Lactate concentration change | -43 [-68, -23]µM*<br>p=0.01* | -90[-200, -27]µM*<br>p=0.06 |
| Lactate base line concentration | -35 [-70, -6]µM*<br>p=0.4 | -80[-140, -42]µM*<br>p=0.03* |

**Table S5**. Difference between ventral and dorsal hippocampi revealed in vivo using [14C]2-deoxy-D-glucose autoradiography in conscious rats

| Publication | Animal, age, condition | Hippocampal structure | Dorsal (µmol/100g/min) | Ventral (µmol/100g/min) |
|---|---|---|---|---|
| Soncrant et al.,1985 (23) | Fischer-344 male rats, 3 months, restrained hind-limbs, T=36.4C | **CA1**<br>CA2<br>CA3<br>DG | **43 ± 3**<br>57 ± 3<br>50 ± 3<br>43 ± 2 | **54 ± 2**<br>58 ± 3<br>58 ± 3<br>52 ± 3 |
| Walovitch et al., 1986 (24) | Fischer-344 male rats, 12-16 months, partially immobilized and placed in a sound-insulated wooden chamber, T=35.1±0.1 °C | **CA1**<br>CA2-CA3<br>DG | **42 ± 3**<br>48 ± 4<br>47 ± 4 | **50 ± 4**<br>53 ± 4<br>50 ± 4 |
| Chastain et al., 1987 (25) | Wistar male rats, 250-300g, free roaming animals temp. not communicated | **CA1** | **48 ± 10** | **61 ± 14** |
| Kelly et al., 1988 (26) | 300-400g, Wistar male rats, restrained hindquarters, temp. not communicated | Subiculum<br>DG | 61 ± 2<br>45 ± 2 | 66 ± 3<br>52 ± 2 |
| Yoshino et al., 1991 (27) | Sprague-Dawley male rats, 250-300g, partially immobilized, T=37-38° | Str. Pyramidale<br>Str. Lacunosum moleculare | **52 ± 3**<br>73 ± 3 | **73 ± 3**<br>83 ± 4 |
| Pereira da Vasconcelos et al., 1992 (28) | Sprague-Dawley rats, post natal day 21, non restricted, T=37°C | All | 38 ± 1 | 39 ± 1 |
